# Long-term labeling and imaging of synaptically-connected neuronal networks *in vivo* using double-deletion-mutant rabies viruses

**DOI:** 10.1101/2021.12.04.471186

**Authors:** Lei Jin, Heather A. Sullivan, Mulangma Zhu, Thomas K. Lavin, Makoto Matsuyama, Xin Fu, Nicholas E. Lea, Ran Xu, YuanYuan Hou, Luca Rutigliani, Maxwell Pruner, Kelsey R. Babcock, Jacque Pak Kan Ip, Ming Hu, Tanya L. Daigle, Hongkui Zeng, Mriganka Sur, Guoping Feng, Ian R. Wickersham

## Abstract

Monosynaptic tracing is a widely-used technique for mapping neural circuitry, but its cytotoxicity has confined it primarily to anatomical applications. Here we present a second-generation system for labeling direct inputs to targeted neuronal populations with minimal toxicity, using double-deletion-mutant rabies viruses. Spread of the viruses requires expression of both deleted viral genes *in trans* in postsynaptic source cells; suppressing this expression with doxycycline following an initial period of viral replication reduces toxicity to postsynaptic cells. Longitudinal two-photon imaging *in vivo* indicated that over 90% of both presynaptic and source cells survived for the full twelve-week course of imaging. *Ex vivo* whole-cell recordings at 5 weeks postinfection showed that the second-generation system perturbs input and source cells much less than does the first-generation system. Finally, two-photon calcium imaging of labeled networks of visual cortex neurons showed that their visual response properties appeared normal for 10 weeks, the longest we followed them.

## Main

Monosynaptic tracing, or the labeling of neurons in direct synaptic contact with a targeted neuronal population using a deletion-mutant neurotropic virus, has become an important technique in neuroscience since its introduction in 2007^1^, because it is the primary available means of brain-wide identification of directly connected neurons. Its core principles are (i) selective infection of a targeted neuronal group of interest with a deletion-mutant neurotropic virus (which in almost all implementations to date is a rabies virus with the envelope glycoprotein gene “G” deleted from its genome) and (ii) complementation of the deletion *in trans* in the targeted starting cells, so that the virus can fully replicate within them and spread, as wild-type rabies virus does, retrogradely (in the central nervous system^2^) across those neurons’ input synapses to neurons directly presynaptic to them. This system has contributed to many significant discoveries about the synaptic organization of many systems in the mammalian nervous system^3–16^.

However, with a few notable exceptions^14–21^, monosynaptic tracing has primarily served simply as an anatomical tool for static identification of connected neurons, because the first-generation, “ΔG” rabies viral vectors that it is based on are swiftly cytotoxic^22–24^. This toxicity stems from the fact that, while deletion of the glycoprotein gene, as intended, prevents the virus from spreading between neurons in the absence of transcomplementation, it leaves the viral transcription and replication machinery intact, so that the virus still rapidly expresses viral genes at high levels and replicates the viral core to high copy numbers^25^, perturbing endogenous gene expression^26, 27^, inhibiting host cell protein synthesis^28^, and killing most infected neurons within approximately two weeks^22, 23^.

Several efforts have been made to engineer less-toxic or nontoxic monosynaptic tracing systems. A first-generation system based on the CVS-N2c strain of rabies virus appears to have lower toxicity than does the widely-used SAD B19 strain^24^. More recently, a paper in *Cell* reported that adding a destabilization domain to the C terminus of the viral nucleoprotein rendered the virus nontoxic, allowing monosynaptic tracing “with no adverse effects on neural physiology, circuit function, and circuit-dependent computations”^29^. We have since shown that those results were probably obtained using reversion mutants that had lost the intended C-terminal addition^30^, although we also showed that the technique may be salvageable^30^, and the authors of the original study have persevered with their approach^31^. From our own laboratory, we have introduced second-generation, “ΔGL” rabies viral vectors, which have the viral polymerase gene “L” (for “large” protein) deleted along with G, and showed that they do not appear to perturb the structure or function of labeled neurons^23^. However, no one has previously shown that these vectors can spread between neurons. Although a baculovirus-based complementation system has recently been reported, it was not shown to work either *in vivo* or otherwise^32^.

Here we show that second-generation (ΔGL) rabies viral vectors can be used for monosynaptic tracing of inputs to genetically-defined populations of neurons, with the double deletion complemented by expression of both deleted viral genes *in trans* in the postsynaptic cells.

## Results

### Construction of a monosynaptic tracing system based on ΔGL rabies viruses

We planned for the second-generation monosynaptic tracing system to use the same pseudotyping strategy^1^ used in the first-generation one for targeting the initial RV infection to the postsynaptic starting cells (or “source cells”, if defined as cells expressing the complementing gene(s) and therefore able to support viral replication). We therefore began by making versions of our previously-introduced second-generation (ΔGL) rabies viral vectors^23^ packaged with the avian retroviral envelope protein EnvA, for selective infection of cells engineered to express EnvA’s receptor, TVA^33, 34^.

Whereas the first-generation system typically relies on an AAV “helper virus” for expression of the deleted glycoprotein gene *in trans* in the source cells^35–38^, the polymerase gene that is also deleted in the second-generation vectors is ∼6.4 kb, too large^39–42^ for a straightforward extension of this approach. We also wanted expression of both the polymerase and the glycoprotein to be regulatable using the tetracycline system^43^, because we anticipated that sustained expression and resulting viral replication could be toxic to source cells.

We therefore generated a knock-in mouse line, “TRE-CB” (short for “TRE-tight-mCardinal-P2A-B19L”), in which the genes for the RV polymerase (SAD B19 strain) and the red fluorophore mCardinal^44^ are under the control of the tetracycline response element in the TIGRE locus^45–47^. To interface with this TRE-driven polymerase allele, we developed a helper virus combination (now published for use with first-generation RV^38, 48, 49^) based on the tet system. Specifically, one AAV, “AAV1-syn-FLEX-splitTVA-EGFP-tTA” is Cre-dependent^50^ (FLEX/DIO design^51^) and expresses TVA, EGFP^52, 53^, and the tetracycline transactivator (tTA), while a second AAV, “AAV1-TREtight-mTagBFP2-B19G”, expresses the RV glycoprotein and the blue fluorophore mTagBFP2^54^ under the control of the tetracycline response element. Our intent was that, when the helper viruses were used in the TRE-CB mouse with Cre expression in starting cells, the tTA expressed by the first AAV would drive expression both of G from the second AAV and of L from the knock-in allele. Note that the use of tTA instead of rtTA makes this a Tet-Off system, so that the genes are expressed by default, with optional administration of a tetracycline analog (e.g., doxycycline) acting to suppress their expression.

ΔGL viruses, by design, express genes at very low levels and are therefore intended to express recombinases, which allow many downstream applications even when expressed at low levels^23^. The viruses in this study therefore encoded either Cre (codon-optimized for mouse^55^) or Flpo (a codon-optimized version of the yeast recombinase Flp^56, 57^). Another necessary component of the system was therefore a means of reporting Cre or Flpo activity, for which we used the Ai14^58^ and Ai65F^47^ mouse lines, in which recombination causes tdTomato expression in cells expressing Cre or Flp, respectively.

Based on our extensive prior experiments titrating these helper viruses for use with first-generation rabies virus to achieve high transsynaptic spread efficiency in Cre mice but low background without Cre^48, 49^, for this second-generation system we used the same helper virus concentrations and seven-day interval between AAV and RV injections as we have previously described for the first-generation one, only varying the survival time after RV injection.

### Labeling inputs to corticostriatal projection neurons: Cre vs Flpo

We targeted corticostriatal cells in primary somatosensory cortex (S1) using retrograde infection by rAAV2-retro^59^ injected into dorsolateral striatum (Fig. 1), with helper AAVs and subsequently RVΔGL injected into S1. In one design (Fig. 1a), the rAAV2-retro expressed Cre, the helper AAVs were Cre-dependent as described above, and the RVΔGL expressed Flpo; these injections were done in the Ai65F reporter line (Flp-dependent tdTomato) crossed to TRE-CB. In the other design (Fig. 1b), the rAAV2-retro expressed Flpo, the helper AAVs were Flpo-dependent (i.e., the first helper virus contained orthogonal FRT sites instead of lox sites), and the RVΔGL expressed Cre instead of Flpo; these injections were done in the Ai14 reporter line (Cre-dependent tdTomato) crossed to TRE-CB. Titers of AAV and RV vectors were matched across the two designs (see Methods). In order to test the effect of using the Tet-Off mechanism to suppress G and L expression after an initial period of unrestricted viral replication, in some conditions we switched the mice to food containing doxycycline two weeks after rabies virus injection (“dox (food)” conditions), while in others (the “dox (inj+food)” conditions) mice also received doxycycline by intraperitoneal injection for three days beginning at the same time as dox food was begun. Two to five weeks after rabies virus injection, mice were perfused and the results examined by confocal microscopy, with transsynaptic spread efficiency quantified by counting labeled neurons in contralateral cortex and ipsilateral thalamus.

**Fig. 1:**
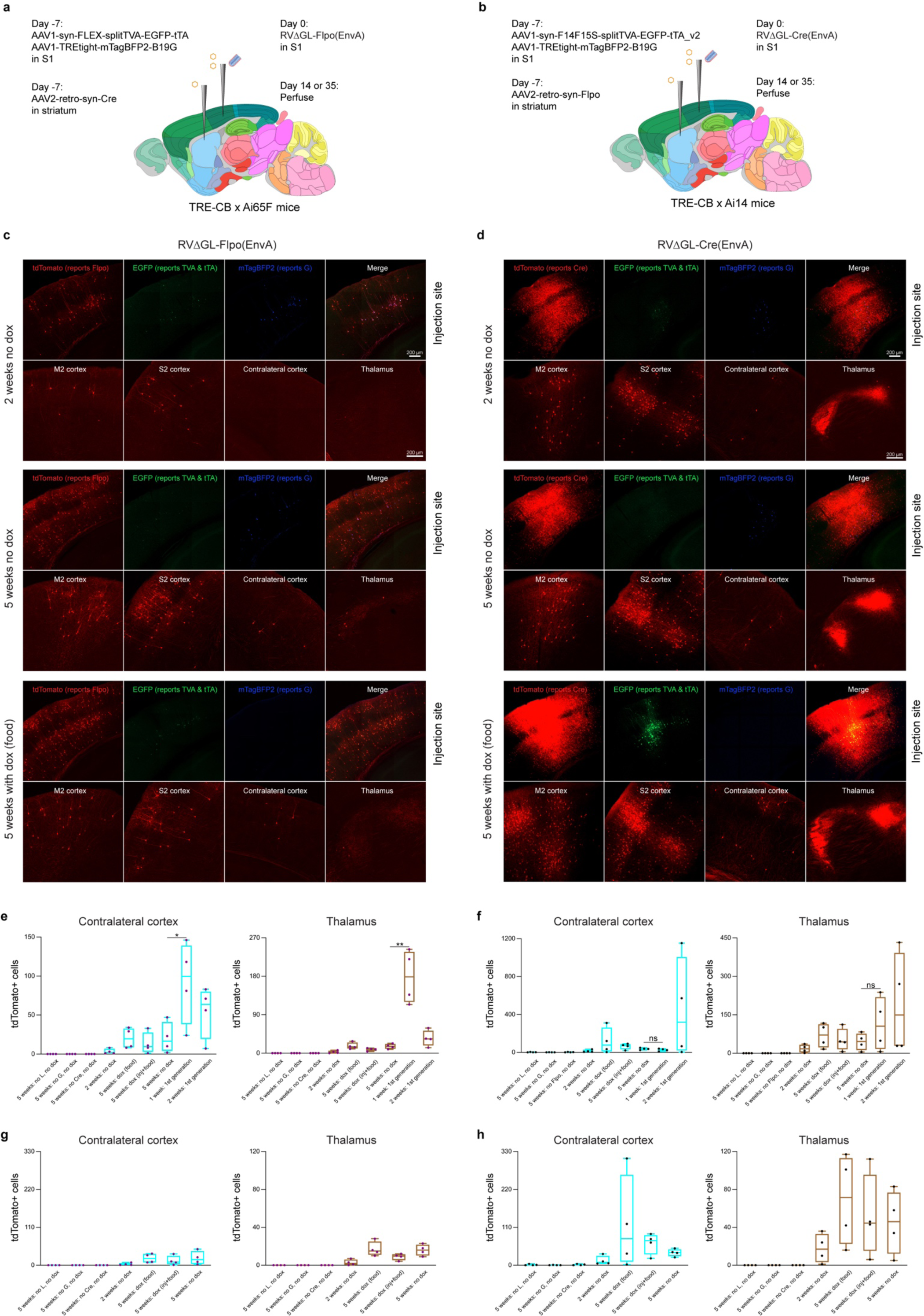
Second-generation monosynaptic tracing of inputs to corticostriatal neurons. **a-b,** Corticostriatal neurons were retrogradely targeted by an AAV2-retro expressing either Cre or Flpo injected in dorsolateral striatum, with a Cre- or Flp-dependent helper virus combination injected in S1. EnvA-enveloped ΔGL rabies virus (RV) expressing either Flpo or Cre was injected in S1 seven days later. The mice were crosses of the “TRE-CB” line described here (providing RV polymerase *in trans*) with tdTomato reporter lines to report activity of the rabies viruses (either Ai65F for Flp or Ai14 for Cre). For the “dox” conditions, mice were switched to doxycycline-containing food 14 days after rabies virus injection, to suppress expression of the viral polymerase and glycoprotein after viral spread; the “dox (inj+food)” mice also received intraperitoneal doxycycline injections for three days beginning 14 days after rabies virus injection. Brain images adapted from the Allen Mouse Brain Reference Atlas (atlas.brain-map.org). **c,** Images of cells labeled with the Flpo-expressing RV. Whereas at 2 weeks few cells were found in input regions, by 5 weeks substantial numbers of labeled cells were found in ipsilateral secondary motor and somatosensory cortices, in contralateral S1, and ipsilateral thalamus, with or without doxycycline (note that doxycycline suppresses expression of mTagBFP2 as well as the viral genes G and L). Considerably fewer input cells were labeled with this version of the second-generation system than with the corresponding first-generation one (panel **e** and Extended Data Fig. S2). Scale bars: 200 µm, apply to all images. **d,** With the Cre-expressing RV and Flp-dependent helper viruses, labeled cells were found in the same regions but in greater numbers. **e-f,** Counts of labeled cells in contralateral S1 and ipsilateral thalamus for RVΔGL-Flpo (**e**) and RVΔGL-Cre (**f**). Counts are the total number of cells found in a series of every sixth 50-µm section spanning each brain, so that the true number of labeled neurons in the entirety of each brain would be approximately six times the number shown here. Note that, although the first-generation Flp-dependent system at two weeks had on average labeled more neurons in input regions than any other condition (**f**), almost no source cells were found in the first-generation conditions at two weeks (see Extended Data Fig. S3), suggesting that almost all source cells had died by that point, consistent with our previous findings regarding the toxicity of first-generation rabies viral vectors. **g-h,** Same data as in **e-f** but with the first-generation counts omitted for clearer display of the ΔGL data and with the ranges of the y-axes of the Flpo and Cre versions set to the same value for each input region for easier comparison of the Flpo vs Cre results. In all experiments with the second-generation system, spread of RVΔGL was greater at five weeks than three weeks, and omission of L, G, or Cre/Flpo resulted in trivial numbers of labeled neurons in input regions (see Extended Data Fig. S2 for counts and representative images for these control conditions). The Flpo-expressing ΔGL RV labeled far fewer cells than did the first-generation version; however, the Cre-expressing ΔGL RV labeled more cells than the Flpo-expressing version in all experimental conditions, although these differences were not statistically significant and were underpowered due to the high variance and low n (=4 mice) for each condition (see Supplementary File S1 for counts and statistical comparisons; see also Extended Data Fig. S4 for example whole-brain image series of mice labeled with each virus combination).

With the Flpo-expressing RVΔGL, when all components were included, there were labeled neurons in the input regions at two different survival times (2 and 5 weeks after RV injection), and doxycycline administration resulted in slightly more spread, though not significantly (Fig. 1c, 1e, and 1g. see Supplementary File S1 for all cell counts and statistical comparisons). This spread of the ΔGL virus to input regions, increasing with time, was despite the fact that we were unable to see clear mCardinal signal, suggesting very low expression of L under the conditions implemented (Extended Data Fig. S1). First-generation control experiments using RVΔG-Flpo with the 7-day survival time that is typical for the first-generation system^48, 60^ labeled significantly more cells in both input regions than any condition using the second-generation RVΔGL-Flpo (Fig. 1e, and see Extended Data Fig. S2 for images from control conditions).

The second-generation system using the RVΔGL expressing Cre instead of Flpo resulted in more labeled neurons in input regions than the one using RVΔGL expressing Flpo (Fig. 1d, 1f, and 1h), although the comparisons between the Cre and Flpo conditions were underpowered due to high variance and modest n (=4 mice per condition throughout our study) and did not achieve statistical significance (see Supplementary File S1 for all cell counts and statistical comparisons).

Control experiments in which either Cre or G was omitted resulted in almost no labeled cells in input regions in either contralateral cortex or thalamus (Fig. 1e-h & Extended Data Fig. S2. See Supplementary File S1 for all cell counts and statistical comparisons). This indicates that the apparent transsynaptic spread was not due to trivial confounding effects, such as direct retrograde infection by the TVA-expressing helper AAV followed by infection of the resulting TVA-expressing axons by the rabies virus, or simply direct retrograde infection by residual RV coated with its native glycoprotein. For more discussion of such possible confounds, see Lavin et al.^48^.

Crucially, control experiments in which L was omitted (by using double transgenics without the TRE-CB allele) also resulted in no label in input regions (Fig. 1e-h & Extended Data Fig. S2). This indicates that complementation with both G and L is required for the ΔGL virus to replicate within the source cells and spread to presynaptic ones. See Supplementary File S1 for all counts and statistical comparisons.

Note that although the first-generation RVΔG-Cre at two weeks postinjection had labeled more input neurons on average than any other condition (**f**), almost no source cells were found at this timepoint with either of the first-generation combinations (none in six out of eight mice, and two cells each in two mice; see Extended Data Fig. S3, panels (**a**) and (**c**), and Supplementary File S1), suggesting that the first-generation system had killed nearly all source cells by that point.

Representative whole-brain image series of labeled inputs to corticostriatal neurons using the Cre-expressing first-and second-generation rabies viral vectors are shown in Extended Data Fig. S4.

### Mapping inputs to dopaminergic and parvalbuminergic neurons in Cre mouse lines

We tested the system in two widely-used Cre mouse lines: DAT-IRES-Cre^61^ and PV-Cre^62^, crossed to TRE-CB and the Flp reporter line Ai65F (Fig. 2 and Extended Data Figs. S5 & S6). In DAT-IRES-Cre, in which Cre is expressed in dopaminergic cells in the midbrain, we injected the viruses in substantia nigra, pars compacta (SNc); in PV-Cre, in which Cre is expressed in parvalbumin-expressing cells in cortex and elsewhere, we injected primary somatosensory cortex (S1).

**Fig. 2:**
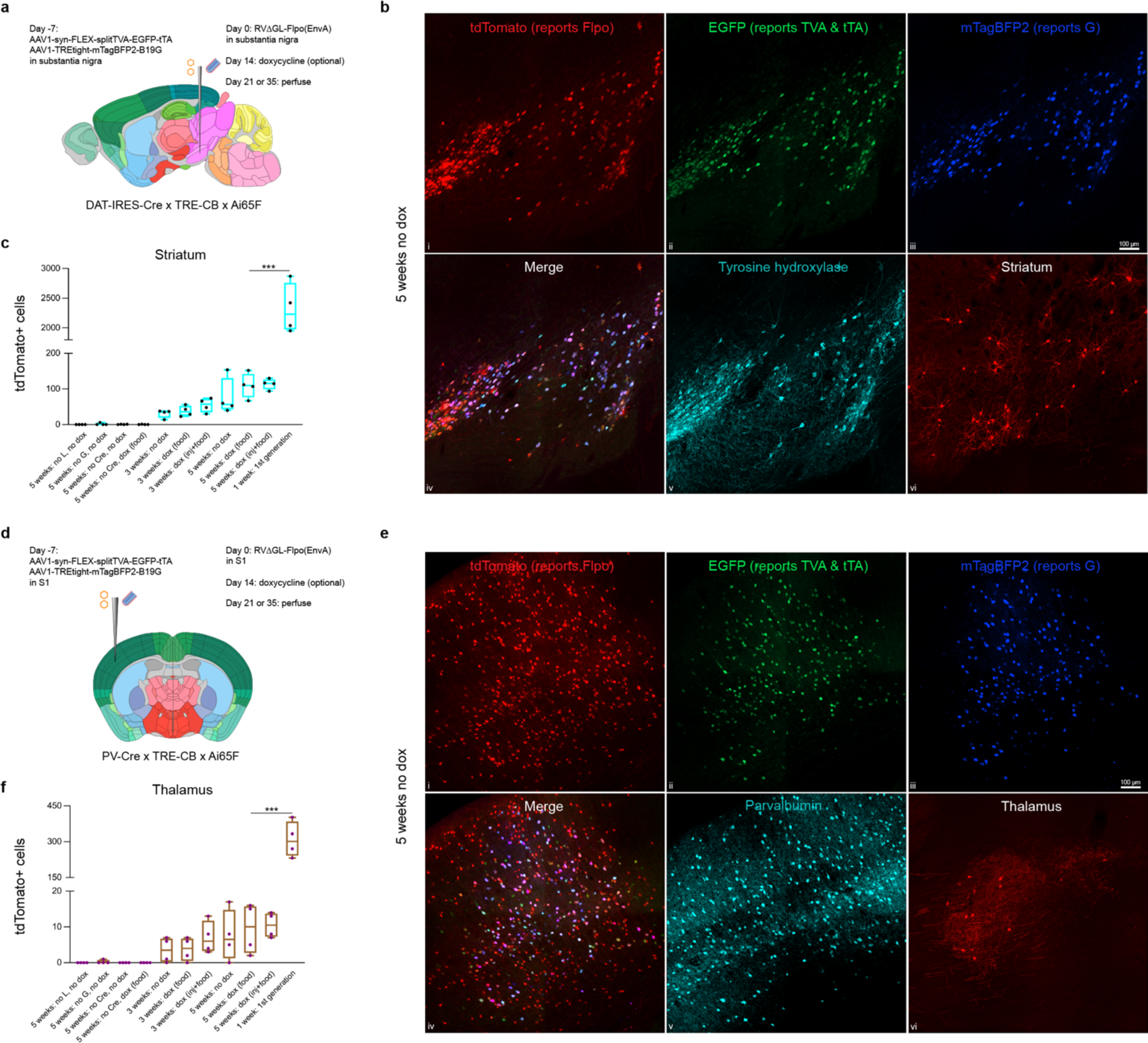
Second-generation monosynaptic tracing in Cre mice. **a,** Experimental design for labeling inputs to dopaminergic cells in the substantia nigra, pars compacta (SNc). The Cre-dependent helper virus combination was injected into the SNc of triple transgenic mice (DAT-IRES-Cre x TRE-CB x Ai65F); EnvA-enveloped ΔGL rabies virus expressing Flpo was injected seven days later. Brain images adapted from the Allen Mouse Brain Reference Atlas (atlas.brain-map.org). **b,** Example of labeled cells five weeks after rabies virus injection, without doxycycline. **b.i-v,** Injection site in SNc, with many cells co-expressing tyrosine hydroxylase (indicating the dopaminergic cells), EGFP (expressed by the first helper virus), mTagBFP2 (expressed by the second helper virus), and tdTomato (reporting activity of the Flpo-expressing rabies virus). Scale bar in **iii**: 100 µm, applies to all images. **b.vi,** Medium spiny neurons in striatum labeled by the ΔGL rabies virus. **c,** Counts of labeled striatal cells (total found in a series of every sixth 50-µm section spanning each brain) in all conditions tested. Omitting any of L, G, or Cre resulted in almost no labeled cells in striatum. Administration of doxycycline to suppress G and L expression after the first two weeks, as well as longer survival time (5 weeks vs 3 weeks), both appeared to result in increased transsynaptic spread, although most comparisons were not statistically significant (see Supplementary File S2 for all counts and statistical comparisons). The first-generation system (with a much shorter survival time) labeled many more cells than all conditions tested for this implementation of the second-generation one. **d,** Experimental design for labeling inputs to parvalbumin-expressing cells in primary somatosensory cortex (S1) in PV-Cre x TRE-CB x Ai65F mice. **e,** Example of labeled cells five weeks after rabies virus injection, without doxycycline. **e.i-v,** Injection site in S1, with many cells coexpressing parvalbumin, reporters of both helper viruses, and tdTomato reporting activity of the rabies virus. In addition to these source cells, many putatively presynaptic cells expressing tdTomato are present. Scale bar in **iii**: 100 µm, applies to all images. **e.vi,** Relay neurons in ipsilateral thalamus labeled by the ΔGL rabies virus. **f,** Counts of labeled thalamic cells (total found in a series of every sixth 50-µm section spanning each brain) in all conditions tested. Again, omitting any of L, G, or Cre resulted in little or no transsynaptic spread, provision of doxycyline appeared to increase transsynaptic spread somewhat, and longer survival time resulted in more spread, although comparisons were underpowered and not statistically significant (see Supplementary File S3 for all statistical comparisons and p-values). The first-generation system again labeled many more cells than did this implementation of the second-generation system, potentially due to low efficiency of recombination by Flpo in these reporter mice.

In both mouse lines, with or without doxycycline, we found spread of rabies virus to regions known to provide input to the respective populations: in DAT-IRES-Cre, we found labeled neurons in striatum (Fig. 2b) and cortex; in PV-Cre, we found labeled neurons in thalamus (Fig. 2e) and other cortical areas, among other locations. In both mouse lines, just as with the corticostriatal experiments described above, there were more neurons at a later time point (five weeks following rabies virus injection) than at an earlier one (three weeks) (see Supplementary Files S2 and S3 for all cell counts and statistical comparisons). Because pilot studies had suggested that survival times of one or two weeks did not result in substantial spread, and that survival times of seven weeks and higher did not result in much more spread than at five weeks, we only compared three- and five-week survival times for this set of quantitative experiments.

A somewhat surprising result from these experiments was that label in input regions in both mouse lines and at both survival times was somewhat higher when doxycycline was administered beginning two weeks after rabies virus injection, although these comparisons were again underpowered and not statistically significant. This was contrary to our expectation that this intervention, which was designed to reduce toxicity to source cells by shutting off viral replication after an initial period, could also reduce the efficiency of transsynaptic spread.

Comparison to the first-generation system in the same Cre lines, however, showed that this implementation of the second-generation one appeared to be far less efficient: matched experiments using a first-generation virus, RVΔG-Flpo (and in mice without the TRE-CB allele), with a seven-day survival time, labeled 1.3-1.5 orders of magnitude (∼21x in DAT-IRES-Cre, ∼33x in PV-Cre) more neurons in input regions (Fig. 2c & 2f).

The results of the control experiments with both mouse lines were as expected: omission of any of G, L, or Cre resulted in almost no labeled cells in input regions (Fig. 2 & Extended Data Figs. S5 & S6; see Supplementary Files S2 and S3 for all counts and statistical comparisons).

### Mapping inputs to dopaminergic neurons in a Flpo mouse line

Because the Flp-dependent version of the system using RVΔGL-Cre resulted in more labeled input neurons than the Cre-dependent one using RVΔGL-Flpo in the corticostriatal experiments (Fig. 1), we tested the Flp-dependent system in DAT-P2A-Flpo mice (crossed to TRE-CB and Ai14), as shown in Fig. 3. We again found the expected patterns of label at the injection site and in the striatum and other input regions, and controls in which either G or Flpo were omitted resulted in very few (0-5) labeled cells in striatum (see Supplementary File S4 for counts and statistical comparisons). Without doxycycline, the numbers of labeled striatal cells were higher with the Flp-dependent system in this mouse line than with the corresponding Cre-dependent condition in DAT-IRES-Cre mice, although this difference was not statistically significant (p=0.134); with doxycycline, however, the numbers were lower than in the corresponding Cre-dependent condition, a difference which was significant. The first-generation system again labeled more striatal cells than did the second-generation system, although in this case the difference was much smaller (mostly because the efficiency of this Flp-dependent implementation of the first generation system was significantly less than that of the Cre-dependent one) and the comparison was not quite statistically significant (p=0.051). See also Extended Data Fig. S7 for whole-brain image series for RVΔG-Cre and RVΔGL-Cre, and see Extended Data Fig. S8 for higher-resolution images of label in several input regions (amygdala, external globus pallidus, and nucleus accumbens) for both ΔG and ΔGL viruses expressing Flpo and Cre.

**Fig. 3:**
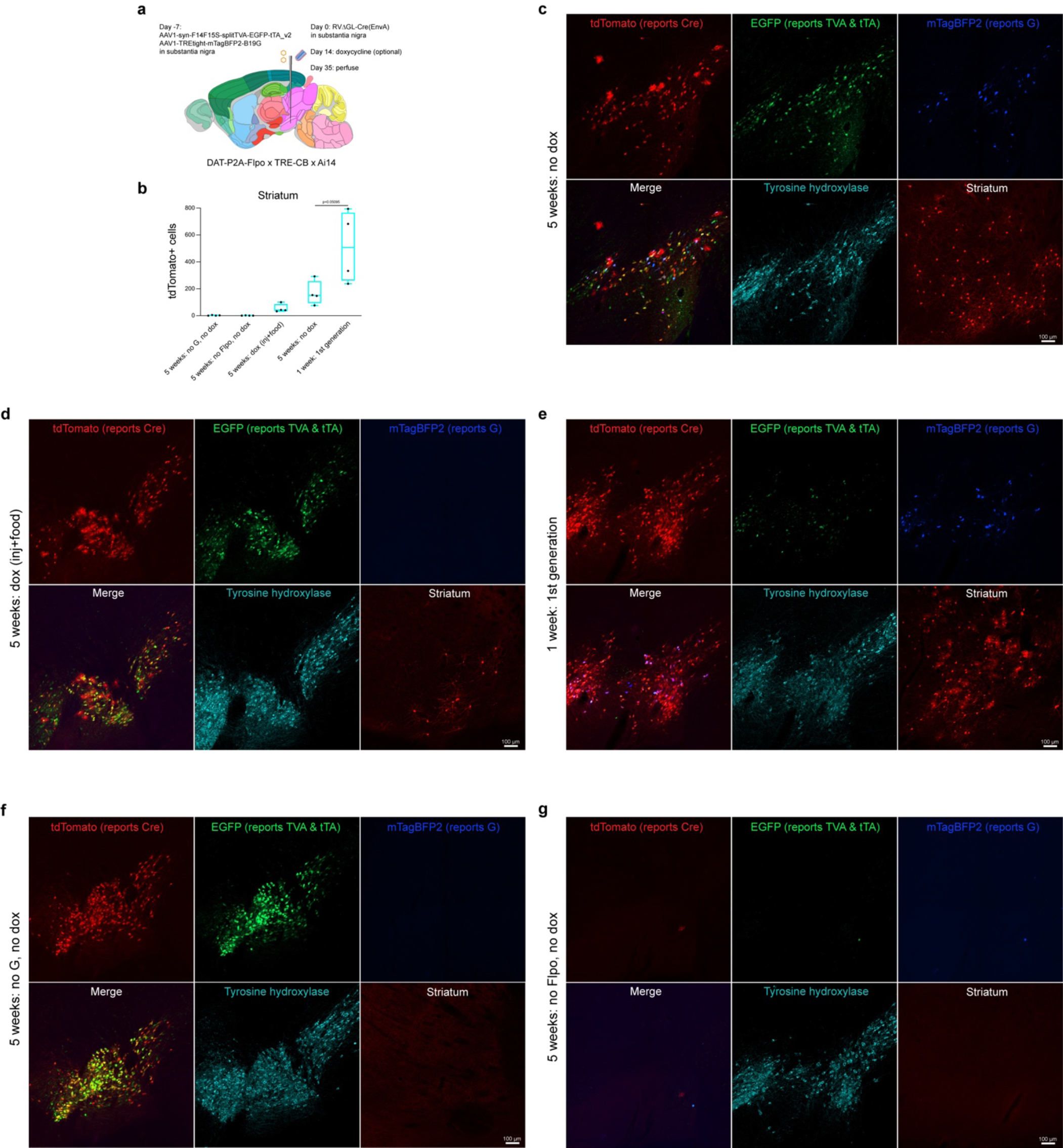
Second-generation monosynaptic tracing in DAT-P2A-Flpo mice. **a,** The design of the experiment was similar to that shown in Fig. 2a but using the Flp-dependent helper virus combination, and Cre-expressing rabies virus, in DAT-P2A-Flpo x TRE-CB x Ai14 mice. Brain image adapted from the Allen Mouse Brain Reference Atlas (atlas.brain-map.org). **b,** Counts of labeled striatal neurons (total found in a series of every sixth 50-µm section spanning each brain) in all conditions tested. **c,** Example images of labeled cells five weeks after rabies virus injection, without doxycycline. Many cells at the injection site in SNc were found to co-express tyrosine hydroxylase and the fluorophores expressed by the helper viruses, as well as tdTomato, reporting activity of the Cre-expressing rabies virus. Medium spiny neurons in striatum were labeled by the ΔGL rabies virus. **d,** Example of a doxycycline (injection + food) mouse; numbers of striatal input neurons were lower in this condition than without doxycycline, although comparisons were again underpowered and not statistically significant. **e,** Example label from the first-generation system. More cells were labeled on average than in the second-generation conditions, although again the differences were not statistically significant. **f-g,** Example of control conditions in which either G (**f**) or Flpo (**g**) was omitted. Scale bars: 100 µm, apply to all images.

### Longitudinal two-photon imaging of labeled networks

We conducted several experiments to assay the survival and physiological status of neurons labeled with the second-generation monosynaptic tracing system. In previous work, we used longitudinal two-photon microscopy to show that neurons labeled with a second-generation (ΔGL) RV stayed alive for as long as we imaged them (in stark contrast to neurons labeled with a first-generation (ΔG) RV) and also that it did not appear to perturb their visual response properties^23^. Because we had only used the Cre-expressing versions of ΔG and ΔGL viruses for the longitudinal two-photon imaging work from our previous publications^23, 30^, for the two-photon work in the present paper we began by conducting a similar set of experiments using the Flpo-expressing RVΔGL to confirm that it was similarly nontoxic (Extended Data Fig. S11). We found that it was: in contrast to the first-generation version, RVΔG-Flpo, which killed over 96% of labeled neurons by 4 weeks postinjection, the second-generation RVΔGL-Flpo left nearly all labeled neurons alive for as long as we imaged them (16 weeks) (specifically, 85 of the 86 neurons seen at 4 weeks postinjection, when the numbers of visibly ΔGL-Flpo-labeled neurons had stabilized – cf. our similar findings with ΔGL-Cre in Chatterjee et al.^23^ – were still found at 16 weeks in the two mice in which we were able to image that long. See Supplementary File S5 for counts). This confirmed that the Flpo-expressing and Cre-expressing ΔGL viruses are similarly nontoxic to labeled cells, consistent with our expectation that, regardless of the transgene, the large reduction in gene expression level caused by deletion of L would eliminate the cytotoxicity caused by the high gene expression and intracellular replication of wild-type and ΔG rabies viruses. These findings suggested that second-generation monosynaptic tracing should be quite nonperturbative to labeled presynaptic neurons (which are only infected with the ΔGL viruses and not the helper viruses), except perhaps to the extent that toxicity to the postsynaptic source cells might perturb cells presynaptic to them by trophic or other network effects.

We then conducted an imaging experiment to assay how toxic the full second-generation monosynaptic tracing system was to neurons, including the source cells, in which G and L were provided (Fig. 4). Similar to the experiments presented in Fig. 2d-f, we injected the Cre-dependent helper virus combination into visual cortex of PV-Cre x TRE-CB x Ai65F mice, followed by RVΔGL-Flpo(EnvA) seven days later. For this set of experiments, we additionally implanted a headmount and optical window (see Methods), and we imaged fields of view at or near the injection sites on a two-photon microscope repeatedly over 14 consecutive sessions beginning at two days postinjection and ending at 12 weeks (Fig 4a). Just as for the previous experiments, half of the mice were given doxycycline beginning at two weeks postinjection.

**Fig. 4:**
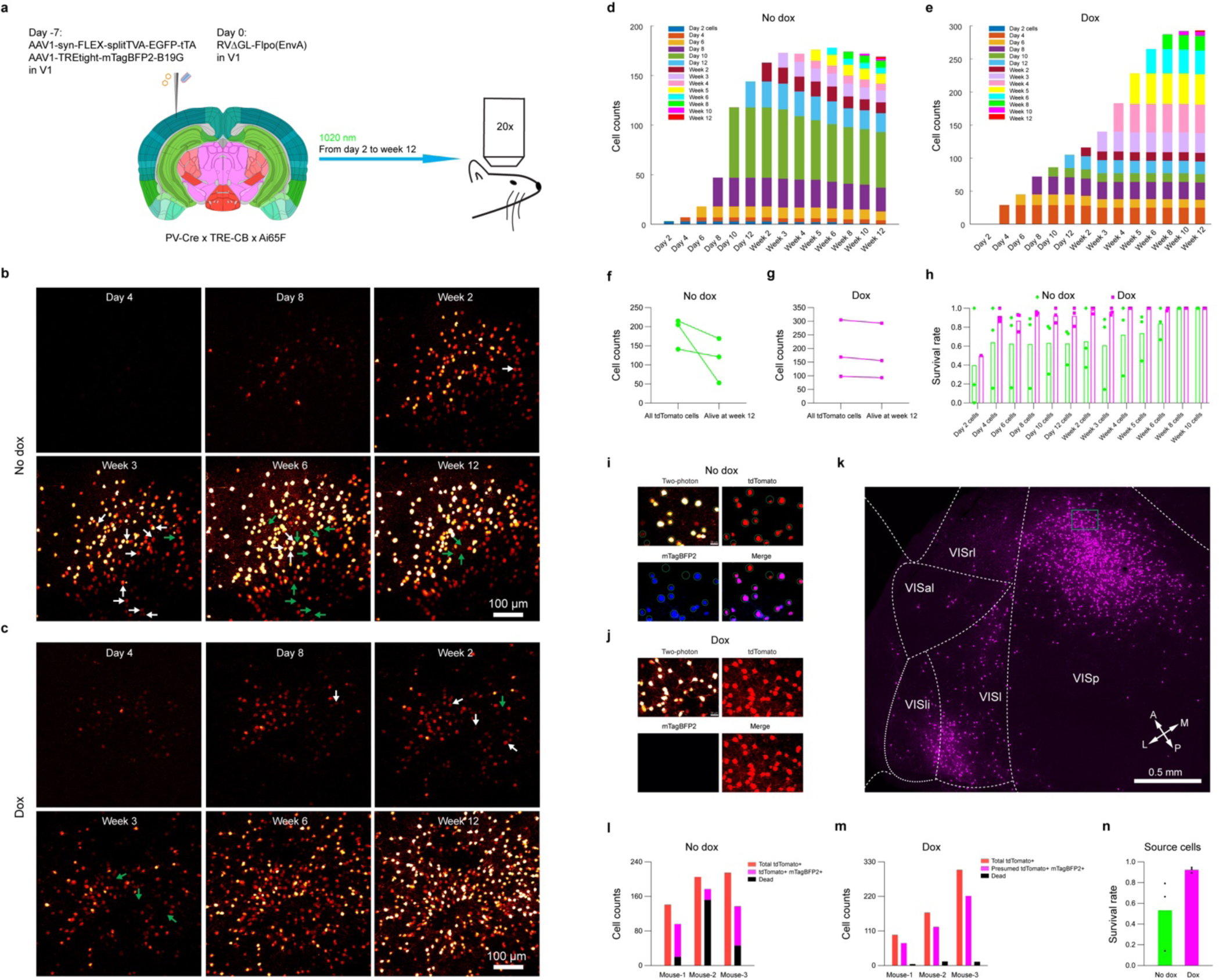
Longitudinal structural two-photon imaging *in vivo*: second-generation monosynaptic tracing preserves >90% of postsynaptic source cells. **a,** Schematic of virus injection and longitudinal two-photon imaging. The helper AAVs, followed by RVΔGL-Flpo(EnvA), were injected in primary visual cortex of PV-Cre x TRE-CB x Ai65F mice, then the injection site was imaged on a two-photon microscope repeatedly for 12 weeks after RV injection. Brain image adapted from the Allen Mouse Brain Reference Atlas (atlas.brain-map.org). b-c, Representative two-photon images at different timepoints of tdTomato-labelled neurons in the injection site in V1. The ‘No dox’ group (b) was fed with regular food throughout, whereas the ‘Dox’ group (c) was fed with doxycycline food (200 mg/kg) starting at two weeks after RV injection until perfusion at week 12, and also received intraperitoneal injections of 100 mg/kg doxycycline every 12 hours for three days, beginning two weeks after RV injection, in order to suppress rabies viral polymerase and glycoprotein expression. Arrows indicate example cells that are alive up to one timepoint (white arrows) but that are missing subsequently (green arrows). In this example, a total of 46 out of 215 cells were lost in the ‘No dox’ group, whereas 12 out of 305 cells were lost in the ‘Dox’ group. Scale bar = 100 μm. d, Cell counts from all 14 timepoints in one of the ‘No dox’ mice. This is the same mouse as shown in b. Bar height for each color indicates the number of cells that were first visible at the corresponding timepoint that are still visible at the timepoint shown on the X axis; for example, the dark green bars indicate the number of cells that were first visible on day 10 that are identifiable at each timepoint from 10 days onward. e, Cell counts from all 14 timepoints in one of the ‘Dox’ mice. This is the same mouse as shown in c. f, Total number of imaged cells and number of cells present at the last imaging timepoint for each of the ‘No dox’ mice. g, Total number of imaged cells and number of cells present at the last imaging timepoint for each of the ‘Dox’ mice. Administration of doxycycline greatly reduces cell attrition. h, Survival rates of cells that appeared at each time point for each mouse in both ‘Dox’ and ‘No dox’ groups. Each data point represents the fraction of cells that first became visible at that timepoint that were still present at the last (12-week) timepoint. i, Aligned 12-week two-photon image (top left) and postmortem confocal images (tdTomato, mTagBFP2, and merged) of an FOV from a ‘No dox’ mouse (the same mouse as used for panel b). Cells found by confocal imaging to have expressed both tdTomato (reporting rabies virus activity) and mTagBFP2 (reporting G expression) were considered to be source cells, along with the cells that died during the 12 weeks of two-photon imaging. Scale bar: 20 µm. j, Aligned 12-week two-photon image (left) and postmortem confocal image of an FOV from a ‘Dox’ mouse (the same mouse as used for panel c). Because doxycycline shut off mTagBFP2 expression along with expression of G and L, it was not possible for us to directly determine the surviving number of source cells in ‘Dox’ mice; for this group, we used an indirect estimate, as shown in panel m below. Scale bar: 20 µm. k, Large-scale confocal image of labeled neurons in multiple visual cortical areas in the same ‘Dox’ mouse. The green rectangle indicates the FOV shown in j. This demonstrates that the transsynaptic spread of ΔGL virus was intact in the mice used for these imaging experiments, just as in the mice used for the similar experiments shown in previous figures. l, Counts for each of the ‘No dox’ mice of the total number of imaged tdTomato+ cells (red), the number of cells that disappeared (black), and the number of cells found by postmortem confocal imaging to have expressed both tdTomato and mTagBFP2 and which therefore were considered surviving source cells (magenta). The fraction of surviving source cells for each of these ‘No dox’ mice was calculated as the ratio (magenta)/(black + magenta). m, Counts for each of the ‘Dox’ mice of the total number of imaged tdTomato+ cells (red), the number of cells that disappeared (black), and the *assumed number of total source cells* (magenta) based on the assumption that the same fraction of total tdTomato+ cells were source cells in the ‘Dox’ mice as in the otherwise-identical ‘No dox’ mice. The fraction of surviving source cells for each of these ‘Dox’ mice was estimated as the ratio (magenta - black)/(magenta). All counts and calculations used for estimating source cell survival rates are given in Supplementary File S6. n, Estimated rates of source cell survival for ‘Dox’ and ‘No dox’ groups. Administration of doxycycline beginning at two weeks postinjection to suppress expression of the two deleted viral genes markedly increases cell survival rate from 53.2% (‘No dox‘) to 92.3% (‘Dox’).

We found that, with or without doxycycline, the majority of tdTomato-labeled cells at the injection sites survived for the full 12 weeks of imaging in most mice. For the mice that did not receive doxycycline, and that therefore had G and L expressing throughout the 12-week experiment, 63.4% of labeled cells survived through the final imaging session. For the mice that did receive doxycycline, however, 94.4% of labeled neurons survived (Fig. 4b-h; see also Extended Data Fig. S12; see also Supplementary File S6 for counts of total and surviving cells). Again, despite doxycycline administration, transsynaptic spread to other cortical areas was still widespread in these mice (Fig. 4k).

Although the overall cell survival rate was high, we anticipated that it would be lower for source neurons, in which G and L are provided and viral replication is therefore restored. In order to determine the survival rate of source cells, the ideal approach would have been to image both tdTomato (reporting activity of the Flpo expressed by the rabies virus) and mTagBFP2 (coexpressed with G by the second helper virus): neurons that expressed both fluorophores would be both infected by the RVΔGL and expressing G (and presumably L, because both depend on tTA expression) and therefore competent hosts for the RV to replicate within and spread to cells presynaptic to them. However, because the two-photon microscope to which we had access was unfortunately incapable of producing short-wavelength light at sufficient power to allow imaging of mTagBFP2 *in vivo*, we had to rely on indirect measures to estimate the source cells’ survival rates. Note that EGFP, which was coexpressed with TVA, is not a good indication of whether a cell is a source cell, because, although some of the cells expressing TVA would have been directly infected by the EnvA-enveloped rabies virus, only a subset of such cells also express G (as reported by mTagBFP2: see Supplementary File S3) and L and are therefore able to support replication and spread of the rabies virus; in any case, at the helper virus concentrations used^48, 49^, the intrinsic EGFP signal is quite weak and needs to be amplified with immunostaining in order to be clearly discernible.

Following longitudinal two-photon imaging of tdTomato-labeled cells over the full 12 weeks, therefore, we perfused the mice, sectioned the brains in a plane approximately parallel to the imaging plane, then imaged the same regions using confocal microscopy. This allowed us to image tdTomato as well as mTagBFP2 (for animals that had not received doxycycline), then align the confocal images with the two-photon ones from the 12-week timepoint to identify the surviving double-labeled source cells (Fig. 4i-j). For those animals that did receive doxycycline, the blue channel contained little signal, as expected (Fig. 4j); our second goal for the confocal imaging, however, was to confirm that transsynaptic spread in these mice used for two-photon imaging had occurred, which it had (Fig. 4k). For both dox and no-dox conditions, we made the assumption that all cells that had died were source cells, based on our finding that cells infected solely with RVΔGL-Flpo without complementation almost always survive (Extended Data Fig. S11; see also Chatterjee et al.^23^ for extensive characterization of the nontoxicity of ΔGL RV).

In the mice that did not receive doxycycline, in which the surviving source cells (i.e., those expressing red and blue fluorophores) were visible by confocal microscopy (Fig. 4i), we estimated the source cell survival rate in each mouse as the ratio of the number of cells in the field of view expressing both red and blue fluorophores divided by the sum of that number plus the number of cells that died in that field of view (Fig. 4l & 4n; see Supplementary File S6 for cell counts and calculations). For these no-doxycycline mice, we obtained a survival rate of 53.2% of source cells.

In the mice that did receive doxycycline, in which surviving source cells were not identifiable as such by postmortem confocal microscopy because they no longer expressed mTagBFP2 (Fig. 4j), we made the assumption that source cells made up the same percentage of total tdTomato-labeled neurons in the fields of view in the dox mice as in the no-dox ones, because the experimental conditions were identical apart from the switch to doxycycline two weeks after RV injection. This gave us an estimated number of total source cells for each mouse; the difference between that and the (small) number of neurons that had disappeared over the course of the two-photon imaging gave us an estimated number of surviving source cells. The ratio of the estimated number of surviving source cells to the estimated total number of source cells was our estimate of the source cell survival rate (Fig. 4m & 4n; see Supplementary File S6 for cell counts and calculations). For these doxycycline mice, we obtained an estimated survival rate of 92.3% of source cells.

### *Ex vivo* whole-cell recordings from labeled input and source neurons

In order to further assay the degree to which the second-generation monosynaptic tracing system perturbs input and source neurons, we performed whole-cell recordings on neurons in acute brain slices made from mice in which inputs to parvalbumin-expressing cortical neurons had been labeled (Fig. 5). As with the anatomical experiments described previously (Fig. 2), we injected Cre-dependent helper viruses in primary sensory cortex of PV-Cre x TRE-CB x Ai65F mice, followed by either RVΔG-Cre(EnvA) or RVΔGL-Cre(EnvA) (or no rabies virus, for AAV-only controls) seven days later. Beginning 35 days after rabies virus injection, we made slices from the injected brains and conducted whole-cell recordings from labeled input cells and source cells to measure their membrane properties.

**Fig. 5:**
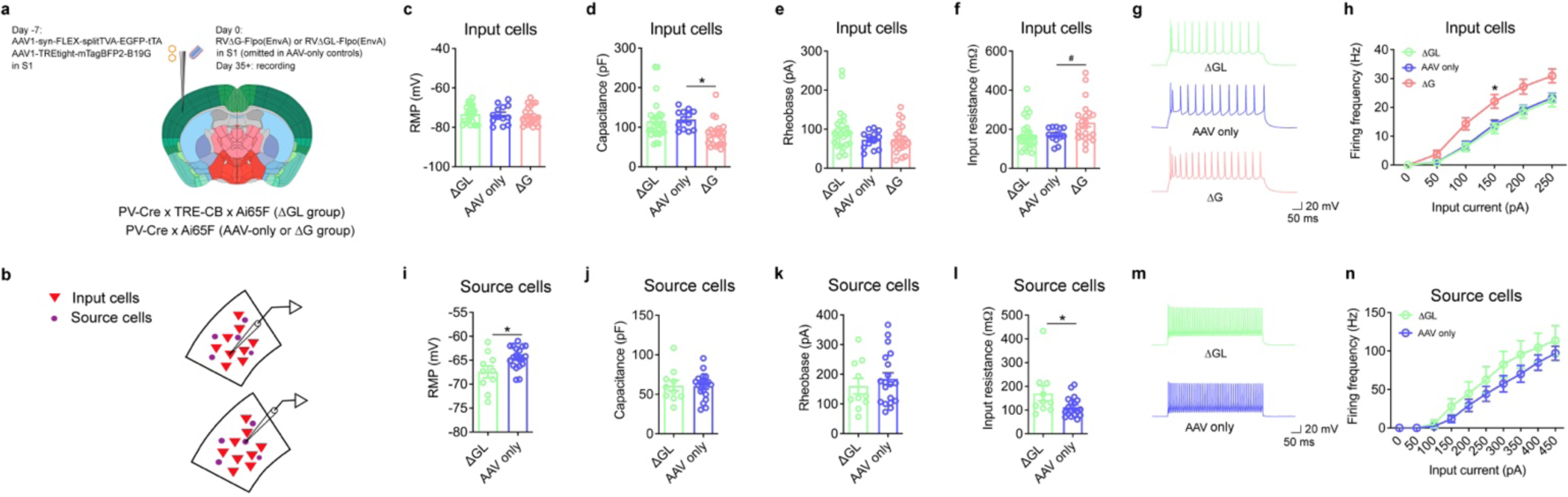
Membrane properties of source cells and input cells labeled by first- and second-generation monosynaptic tracing. **a-b,** Experimental design. Cre-dependent helper viruses were injected in S1 of either PV-Cre x Ai65F x TRE-CB het/het/het (ΔGL group) or PV-Cre x Ai65F het/het (ΔG and AAV-only groups); 7 days later, either RVΔGL-Flpo(EnvA) or RVΔG-Flpo(EnvA) was injected, with an AAV-only control group receiving no rabies virus injection. Beginning 35 days after RV injection and continuing for up to 10 days later, brain slices were prepared, and labeled neurons (“input cells” defined as pyramidal neurons expressing tdTomato alone, or “source cells” defined by neurons expressing both tdTomato and mTagBFP2) were targeted for whole-cell recordings. For AAV-only mice, “source cells” denotes neurons expressing mTagBFP2, and “input cells” denotes randomly-selected unlabeled pyramidal neurons nearby. Brain image adapted from the Allen Mouse Brain Reference Atlas (atlas.brain-map.org). **c-h,** Membrane properties of input cells (as defined above) in the three experimental groups. **c-f,** Resting membrane potential (RMP), membrane capacitance (Cm), rheobase, and input resistance (Rm) of input neurons. With the second-generation system, all four of the measured membrane properties remain normal (not statistically different from the corresponding AAV-only control measurements) 5 weeks after injection of ΔGL rabies virus. With the first-generation system, the few labeled neurons found at this long timepoint had capacitance and input resistance significantly different from controls. ΔGL, 26 cells from 4 mice; AAV control, 13 cells from 3 mice; ΔG, 22 cells from 3 mice; RMP, one-way ANOVA, F(2, 58) = 0.35, p = 0.71. Capacitance, one-way ANOVA, F(2, 58) = 4.924, p = 0.011, Dunnett’s multiple comparisons test, ΔGL vs. AAV control, p = 0.93, ΔG vs. AAV control, p = 0.023, *p < 0.05. Rheobase, one-way ANOVA, F(2, 58) = 1.81, p = 0.17. Input resistance, one-way ANOVA, F(2, 58) = 4.13, p = 0.021, Dunnett’s multiple comparisons test, ΔGL vs. AAV control, p > 0.99, ΔG vs. AAV control, #p = 0.060. **g,** Representative traces of the action potential firing of input cells in response to 250 pA current injection. **h,** Plot of action potential firing frequency in response to sequential current step injection showed increased and normal excitability in the principal neurons labeled by ΔG and ΔGL rabies virus, respectively. Two-Way ANOVA, F(2, 52) = 5.61, p = 0.0062. Dunnett’s multiple comparisons test, ΔGL vs. AAV control, p = 0.94, ΔG vs. AAV control, p = 0.031, *p < 0.05. **i-n,** Membrane properties of source cells (as defined above) in the ΔGL and AAV-only groups. Note that no surviving source cells could be found in the ΔG group at this long (5-week) survival time. **i-l,** Resting membrane potential (RMP), membrane capacitance (Cm), rheobase, and input resistance (Rm) of source cells (as defined above) in the two groups (ΔGL and AAV-only) in which extant source cells could be found. Source cells in the ΔGL group showed more negative resting membrane potential and higher input resistance. AAV control, 19 cells from 3 mice, ΔGL, 10 cells from 4 mice). RMP, unpaired t-tests (two-tailed), p = 0.016, *p < 0.05. Capacitance, unpaired t-tests (two-tailed), p = 0.95. Rheobase, unpaired t-tests (two-tailed), p = 0.47. Input resistance, unpaired t-tests (two-tailed), p = 0.040, *p < 0.05. **m,** Representative traces of the action potential firing of source cells in response to 450 pA current injection. **n,** Action potential firing frequency in response to sequential current step injection showed similar excitability in the source cells of the ΔGL and AAV-only groups. Two-way ANOVA, F(1, 27) = 1.42, p = 0.24.

In brains labeled by the second-generation system, we found that the membrane properties of input cells did not differ from those of unlabeled neurons in the AAV-only control mice, and that source cells differed from “source cells” in the AAV-only control mice (i.e., cells labeled by the helper AAVs) in two of the five measured membrane properties.

By contrast, in brains labeled by the first-generation system, those input cells that remained at the time of recording (5 weeks after infection) differed from unlabeled control neurons in three out of five measured membrane properties (cf. Chatterjee et al.^23^), and no remaining source cells could be found at all. This marked cytotoxicity of the first-generation system to source cells after 5 weeks is unsurprising but illustrates a major difference between the first- and second-generation systems.

### Longitudinal two-photon calcium imaging of cortical neurons’ responses to visual stimuli

Finally, we conducted experiments designed to provide proof of concept for long-term functional imaging of synaptically-connected networks of neurons (Fig. 6). We began by making a Flp-dependent jGCaMP7s reporter AAV and conducted pilot studies in PV-Cre mice to determine a dilution that would not itself cause mass cytotoxicity due to overexpression of jGCaMP7s^47, 63^ (Extended Data Fig. S13). We then conducted an experiment similar in design to the structural imaging study, except that the rabies virus was mixed with the jGCaMP7s AAV before injection and headmount/coverslip implantation (Fig. 6a). We then imaged labeled neurons at or near the injection sites over a series of sessions spanning ten weeks. We imaged the jGCaMP7s signal while the awake mice viewed a series of drifting grating stimuli (Fig. 6b-c); we also imaged tdTomato fluorescence in the same fields of view to confirm that jGCaMP7s expression was restricted to RV-labeled neurons (Fig. 6c). As shown in Fig. 6d-h, as well as Extended Data Figs. S13 and S15, we observed clear orientation tuning curves in many of the labeled neurons, with visual responses appearing to remain stable across repeated imaging sessions over several months (Fig 6d-h, and Extended Data Fig. S15). Following the full course of two-photon imaging, we conducted postmortem sectioning and confocal imaging to confirm that viral spread had occurred in the imaged mice (Extended Data Fig. S16).

**Fig. 6:**
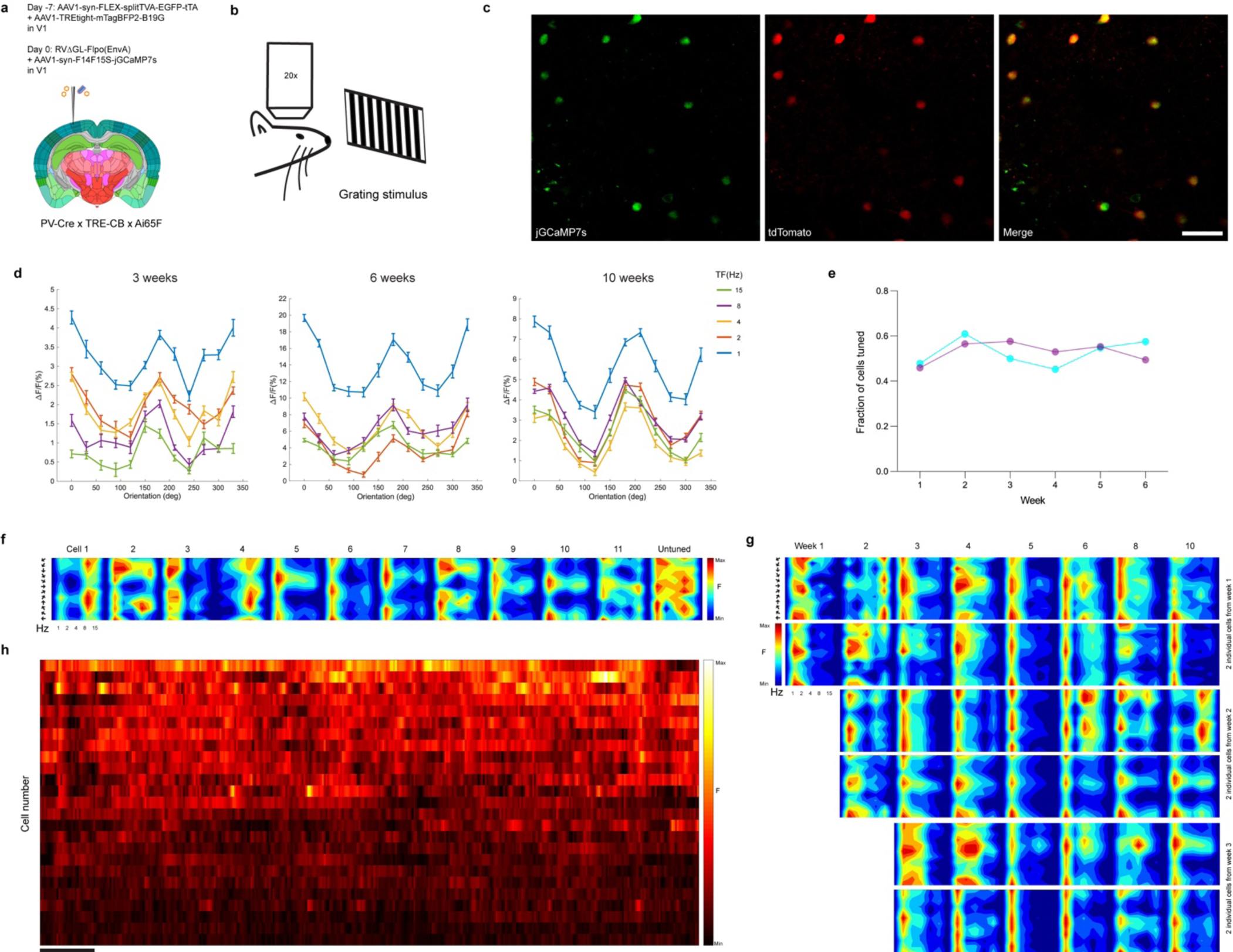
Functional two-photon imaging of transsynaptically-labeled neurons’ visual response properties over ten weeks. **a,** The experimental design was similar to that shown in Fig. 4 but with a Flp-dependent jGCaMP7s AAV injected along with the rabies virus. Brain image adapted from the Allen Mouse Brain Reference Atlas (atlas.brain-map.org). **b,** Following the virus injections, the injection site was imaged on a two-photon microscope while the awake mice were presented with drifting grating stimuli of different orientations and temporal frequencies, repeatedly for 10 weeks after RV injection. **c,** Representative two-photon field of view of neurons expressing jGCaMP7s (green channel) and tdTomato (red channel). Scale bar: 50 µm. **d,** Tuning curves of a jGCaMP7s-expressing neuron obtained with drifting gratings presented at 12 directions of motion and 5 temporal frequencies (TF), repeated 10 times (mean Δ F/F ± s.e.m.) at three different time points (left: week 3; middle: week 6; right: week 10). **e,** Fraction of cells tuned at six different timepoints in two mice. The number of tuned neurons does not decline with time, suggesting intact response properties despite labeling by RVΔG. **f,** Tuning patterns at week 10 of 11 jGCaMP7s-expressing neurons showing clear preferred direction tuning, as wells as the tuning pattern of an untuned neuron, representative of roughly half of imaged cells. **g,** Direction tuning patterns of six individual cells recorded at multiple time points (from week 1 to week 10). The top two cells became visible at week one; the middle two appeared at week two, and the bottom two cells appeared at week three. Tuning patterns remain stable over the entire imaging period. **h,** Single-cell fluorescence time courses for 25 cells, showing activity over the first 120 s of visual stimulation. Cells are ranked in descending order of total activity. Scale bar: 10 s.

## Discussion

We have shown here that it is possible to use nontoxic double-deletion-mutant rabies viral vectors to label direct inputs to neurons targeted based on either their extrinsic projections or their gene expression (using Cre or Flp lines). We have shown that use of the Tet-Off system results in relatively little toxicity even to the source cells. Using whole-cell recordings from labeled neurons, we found that input cells’ membrane properties were not significantly different from those of unlabeled control neurons, and source cells were perturbed in two out of four measured membrane properties; this represents a considerable improvement over the first-generation system, which, at the five-week survival time we examined, left input cells perturbed and source cells dead. We have further used two-photon functional imaging of live second-generation monosynaptic tracing *in vivo* to show that apparently normal visual response properties of labeled neurons can be recorded for at least ten weeks.

It is perhaps surprising that so many source cells (over 90%) appeared to survive when doxycycline was administered beginning at two weeks postinjection (Fig. 4). This was not entirely accidental: we designed the system so that the level and time course of L expression could be titrated as necessary to reduce toxicity to source cells. However, it was undoubtedly fortunate that administering doxycycline beginning two weeks after ΔGL rabies virus injection turned out to be sufficient to preserve the majority of source cells (at least for cortical cells in the PV-Cre mice that we tested this in). It was even more fortunate that this intervention did not appear to reduce the efficiency of transsynaptic spread in either of the Cre lines or in the corticostriatal experiments, although these comparisons were underpowered and not statistically significant.

As we have mentioned throughout, many of the comparisons of the efficiency of the various *in vivo* conditions in our study were underpowered, given to the high variance we found in most conditions and the modest number of mice we used per condition (n=4) because of the large numbers of conditions that we tested. For example, for the corticostriatal experiments shown in Fig. 1, comparing the ΔGL-Cre and ΔGL-Flpo at 5 weeks without dox, the mean numbers of labeled cells in contralateral cortex were 36.5 and 20.5, respectively, with a p-value of 0.20 (Supplementary File S1; numbers are totals across every sixth section; see Methods). Given the pooled standard deviation of 22.38, the number of mice needed to adequately power this comparison would have been 31 in each group. Similarly, for the labeled thalamic neurons in the same groups, we found means of 45 with ΔGL-Cre and 16 with ΔGL-Flpo, with a p-value of 0.14. The pooled standard deviation for these counts was 33.86, indicating that 22 mice per group would have been required for adequate power. We speculate that the high variability seen in many of the conditions may be in part due to our injecting the helper viruses and rabies virus in two successive surgeries, as is typical (and in the case of the corticostriatal experiments, a third injection – AAV2-retro in striatum – was made). It may be that simply mixing the helper AAVs and rabies virus and injecting them together (as some have successfully done^64^) would decrease variability, but we have not tested this approach so far.

A drawback of the second-generation system introduced here is its slow time course, necessitating multi-week incubation times (Figs. 1-3). We presume that this slow time course is due to low L expression from the TRE-CB allele given the low level of tTA expressed by the intentionally-dilute Cre- or Flp-dependent helper virus^48, 49^; this hypothesis is consistent with our inability to detect clear mCardinal signal by confocal microscopy (Extended Data Fig. S1) and perhaps also explains the high percentage of surviving source cells that we found. While a slow time course is inconvenient, it may be the necessary price of preserving the source cells.

An interesting finding from the current study was that using an RVΔGL expressing Cre instead of one expressing Flpo resulted in increased numbers of labeled cells under some conditions (Figs. 1-3), although again these comparisons were not statistically significant. Our working hypothesis is that such a difference, if real, could be due to a difference in inefficiency of the two recombinases^57, 65, 66^, with Cre more faithfully reporting the actual spread of the virus than Flpo when the recombinases are expressed at low levels (e.g., by ΔGL rabies viruses). A relative inefficiency of Flpo could also, however, cause lower efficiency of the Flp-dependent helper virus combination itself relative to the Cre-dependent one; this is consistent with the first-generation system’s lower efficiency in the Flp-dependent corticostriatal experiments than in the Cre-dependent ones (Fig. 1), and similarly in DAT-P2A-Flpo mice (Fig. 3) than in DAT-IRES-Cre mice (Fig. 2). In this study we simply matched the titers of the Flp-dependent combination to those used for the Cre-dependent one, but titration and further optimization of the Flp-dependent helper virus might improve performance.

We note that, even apart from the rabies virus and its complementing gene products, several of the components of the system have the potential to cause cytotoxicity, including tTA^47, 67, 68^, GCaMPs^47, 63^, and even AAV itself^69^. These can be expected to contribute to the toxicity to source cells, perhaps making it more remarkable that we found that so many of them survived, albeit with some perturbation to their membrane properties (Fig. 5). The toxicity to source cells may also depend on the cell type, both because neuron types can be differentially vulnerable to adverse conditions^70, 71^ and because differential tropism for AAV1^49^ (or whatever other serotype might be used for the helper viruses) will change the dosage of tTA, G, L, TVA (and therefore the initial multiplicity of infection by the injected RV), and the fluorophores.

The presynaptic cells, on the other hand, are less burdened than the postsynaptic ones: because we have found that ΔGL RV by itself (in combination with tdTomato expression in Ai14 or Ai65F mice) does not kill neurons, the vast majority of labeled presynaptic cells can be expected to survive. It seems possible that residual toxicity to the source cells postsynaptic to them could cause some remodeling or toxicity via retrograde transneuronal effects^72^; however, our electrophysiological recordings did not detect any perturbation of the input cells’ membrane properties.

While the second-generation system is more complicated than the first-generation one in that it requires two additional transgenic alleles (for expressing L and for reporting activity of the recombinase expressed by the RV), in practice this is not overly burdensome: we have found that the TRE-CB mice can be readily maintained as double homozygotes with the Ai14 or Ai65F alleles, so that production of triply (Figs. 2-6) or doubly (Fig. 1) heterozygous mice is achieved by a single cross with either a recombinase-expressing (e.g., PV-Cre) or wild-type mouse.

Perhaps importantly, our finding that ΔGL rabies viruses do not spread when not complemented by L expression *in trans*, even when G is provided *in trans*, indicates that deletion of either L or G alone is sufficient to prevent transsynaptic spread. This suggests the possibility of a third-generation monosynaptic tracing system, based on single-deletion-mutant ΔL viruses^73^ complemented by expression *in trans* of just L. Because of the reduced complexity of complementing with (once again) only a single gene, such a system may have advantages over the second-generation one described here.

## Supporting information

Extended Data Fig. S4: Series of whole-brain images of labeled inputs to corticostriatal neurons using dG and dGL RV vectors expressing Cre

Extended Data Fig. S7: Whole-hemisphere image series of labeled inputs to dopaminergic neurons using dG and dGL RV vectors expressing Cre

Supplementary File S1: Counts and statistics for corticostriatal experiments.

Supplementary File S2: Counts and statistics for DAT-IRES-Cre experiments.

Supplementary File S3: Counts and statistics for PV-Cre experiments.

Supplementary File S4: Counts and statistics for DAT-P2A-Flpo experiments.

Supplementary File S5: Counts for dG-Flpo and dGL-Flpo comparison.

Supplementary File S6: Counts and calculations for estimation of starting cell survival rates.

Supplementary File S7: Counts of tuned and untuned cells in functional imaging experiments.

## Methods

All experiments involving animals were conducted according to NIH guidelines and approved by the MIT Committee for Animal Care. Mice were housed 1-5 per cage under a normal light/dark cycle for all experiments.

### Cloning

pAAV-syn-F14F15S-jGCaMP7s (Addgene 178514) was made by cloning a Flp-dependent FLEX arrangement of mutually-incompatible FRT sites^74^ into pAAV-synP-FLEX-EGFP-B19G (Addgene 59333) followed by the jGCaMP7s^75^ gene from pGP-CMV-jGCaMP7s (Addgene 104463).

pAAV-syn-F14F15S-splitTVA-EGFP-tTA_v2 (Addgene 183352) was made similarly but with the tricistronic open reading frame from pAAV-syn-FLEX-splitTVA-EGFP-tTA (Addgene 100798) except for the use of FRT sites instead of lox ones (and with a 2-bp frameshift of the in-frame FRT site to prevent creation of a premature stop codon). Note that this plasmid is a replacement for pAAV-synP-F14F15S-splitTVA-EGFP-tTA, which was described in an earlier preprint of this manuscript.

pAAV-syn-FLEX-tTA (Addgene 178516) was made by cloning a codon-optimized tet transactivator gene^76^ into pAAV-synP-FLEX-EGFP-B19G (Addgene 59333).

pAAV-syn-Flpo (Addgene 174378) and pAAV-syn-mCre (Addgene 178515) were made by replacing the FLEX cassette from pAAV-synP-FLEX-EGFP-B19G (Addgene 59333) with mouse-codon-optimized genes for Flp^57^ and Cre^55^.

pCAG-hypBase was made by synthesizing a fragment encoding the “hyperactive” piggyBac transposase iPB7^77^ and cloning into the EcoRI & NotI sites of pCAG-GFP^78^ (Addgene 11150).

The piggyBac plasmid pB-CAG-B19L-IRES-mCherry-WPRE-BGHpA (Addgene 178518) was made by cloning the CAG promoter from pCAG-B19G (Addgene 59921), the SAD B19 L gene, the EMCV IRES^79^, the mCherry^80^ gene, and the woodchuck post-transcriptional regulatory element and bovine growth hormone polyadenylation signal from pCSC-SP-PW-GFP (Addgene 12337), into PB-CMV-MCS-EF1-Puro (System Biosciences #PB510B-1).

The “TLoop”-style^81^ lentiviral transfer vectors pLV-TTBG (Addgene 115232) and pLV-TTBL (Addgene 115233) were made by replacing the CMV promoter and EGFP gene in pCSC-SP-PW-GFP (Addgene 12337) with the leaky tetracycline response element from pAAV-FLEX-hGTB (Addgene 26196) followed by a tricistronic open reading frame consisting of the genes for the “improved tetracycline transactivator” itTA^82^, mTagBFP2^54^, and either the glycoprotein or polymerase gene, respectively, of the rabies virus SAD B19 strain, separated by P2A elements.

### Mouse strains

Mice used in this study were crosses (triple transgenic for DAT-IRES-Cre and PV-Cre experiments, double transgenic for corticostriatal experiments, double or single transgenic (either Cre-negative or L-negative) for control experiments) of the following strains, all in a C57BL/6J (Jackson Laboratory 000664) background:

PV-Cre^62^, DAT-IRES-Cre^61^, Ai14^58^, and DAT-P2A-Flpo^83^ were purchased from Jackson Laboratory (catalog #s 017320, 006660, 007914, and 035436).

The Flp-dependent tdTomato reporter line Ai65F was obtained in our case by crossing the Cre- and Flp-dependent tdTomato double-reporter line Ai65D^46^ (Jackson Laboratory 021875) to the Cre deleter line Meox2-Cre^84^ (Jackson Laboratory 003755), then breeding out the Meox2-Cre allele. An equivalent Ai65F line, made using a different Cre deleter line, was described in Daigle et al. ‘18^47^ and is now available from Jackson Laboratory with catalog # 032864).

The L-expressing mouse line TRE-CB (TRE-tight-mCardinal-P2A-B19L, being distributed by Jackson Laboratory with catalog # 036974) was generated by cloning the genes for mCardinal^44^ and the rabies virus (SAD B19) polymerase, separated by a picornavirus 2A element^85^, into “Ai62(TITL-tdT) Flp-in replacement vector” (Addgene 61576), to make the Flp-in construct “TRE-mCardinal-P2A-B19L Flp-in vector” (Addgene # 178519). The lox-stop-lox sequence present in the original Flp-in vector was removed to make the new construct, so that the tetracycline response element TRE-tight^86^ drives mCardinal-P2A-B19L directly with no dependence on Cre recombination. This plasmid was then used for FLP-mediated targeted insertion into the TIGRE locus^45^ in Ai99 embryonic stem cells as described^47^. Clones verified as containing the correct insert were used for production of knock-in mice by the Mouse ES Cell & Transgenic Facility at the Koch Institute for Integrative Cancer Research at MIT. All subsequent breeding steps were with mice of a C57BL/6J background.

### Production of lentiviral vectors for making cell lines

Lentiviral vectors were made as described^87^ but with the VSVG expression vector pMD2.G (Addgene 12259) as the envelope plasmid and with the following transfer vectors:

pLV-TTBL (described above), to make the L-expressing lentiviral vector LV-TTBL(VSVG),

pLV-U-TVA950 (Addgene 115236), to make the TVA-expressing lentiviral vector LV-U-TVA950(VSVG).

### Production of cell lines

To make the L-expressing cell line BHK-B19L, BHK-21 cells (ATCC# CCL-10) were transfected with pCAG-hypBase and pB-CAG-B19L-IRES-mCherry-WPRE-BGHpA; the cells were expanded into two 15c plates, then resuspended and sorted on a BD FACS Aria to collect the brightest 5% of mCherry-expressing cells.

BHK-B19L-TVA950 cells were made by infecting BHK-B19L cells (described above) with LV-U-TVA950(VSVG) at a multiplicity of infection of approximately 100. Cells were then expanded and frozen in medium containing 10% DMSO for subsequent use.

The BHK-EnvA2-TTBL2 cell line, expressing both the “EnvARGCD” fusion protein^1^ and the RV polymerase, was made by infecting BHK-EnvA2 cells^88^ with the L-expressing lentiviral vector LV-TTBL(VSVG) (described above) at a multiplicity of infection of approximately 100. The cells were expanded into three 15 cm plates, then sorted on a BD FACS Aria to collect those cells with the highest 2% of both green (EnvA) and blue (L) fluorescence. After expanding the collected cells again, we observed that blue fluorescence was so dim as to be difficult to detect by eye; we therefore sorted the cells a second time, again retaining cells with the brightest 2% fluorescence in both channels. The twice-sorted cells, now called BHK-EnvA2-TTBL2, were expanded again and frozen in medium containing 10% DMSO for subsequent use.

### Production and titering of adeno-associated viruses

AAV genome plasmids pAAV-syn-FLEX-splitTVA-EGFP-tTA (Addgene 100798) and pAAV-TREtight-mTagBFP2-B19G (Addgene 100799) have been described previously^38^. These genomes were packaged in serotype 1 AAV capsids by Addgene (catalog numbers 52473-AAV1, 100798-AAV1, and 100799-AAV1). The two helper AAVs from Addgene were diluted in Dulbecco’s phosphate-buffered saline (DPBS) (Thermo, 14190-250) to final titers of 7.22E+10 genome counts (gc)/mL for AAV1-syn-FLEX-splitTVA-EGFP-tTA and 6.50E+11 gc/mL for AAV-TREtight-mTagBFP2-B19G, then combined in a 50/50 ratio by volume before injection. AAVs from Addgene were initially thawed and aliquoted in 20x 5ul aliquots; to make working dilutions, a 5ul aliquot of undiluted virus was thawed, diluted in DPBS (Thermo Fisher 14190-250) to the desired working dilutions (1:20 or 1:200)^48, 49^. pAAV-syn-FLEX-tTA (described above) was packaged as serotype 1 by the UNC Vector Core with titer of 2.51E+12 gc/mL.

rAAV2-retro-syn-Cre, rAAV2-retro-syn-Flpo, AAV1-syn-F14F15S-jGCaMP7s, and AAV1-syn-F14F15S-sTpEptTA_v2 were made by transfecting 8×15c plates (per virus) of HEK 293T cells using Xfect transfection reagent (Takara 631318). Briefly, plates were treated with poly-L-lysine solution were transfected with pAAV vector, pHelper (Cellbiolabs VPK-421), and the appropriate rep/cap plasmid (“rAAV2-retro helper” (Addgene 81070) or pAAV-RC1 (Cellbiolabs VPK-421)). 4 hours after transfection, the medium from plates was aspirated and replaced with 15mL DMEM (Thermo #11995-081) containing 2% fetal bovine serum (FBS) (HyClone SH30071.02) and antibiotic/antimycotic (Thermo Fisher 15240-062). 72 hours after transfection, the supernatants were collected from each plate and replaced again with 15mL DMEM with 2% FBS and antibiotic/antimycotic. Supernatants were stored in 50mL tubes at 4°C. 120 hours after transfection, cells and supernatants were collected and transferred into sterile 50mL tubes. Cells were pelleted by centrifugation and washed with DPBS, and supernatants were transferred into a sterile bottle for PEG precipitation. Cells were lysed by four freeze/thaw cycles and stored at −80°C until final thaw and purification steps. 50mL of cold 40% polyethylene glycol 8000 (PEG) in 2.5M NaCl was added to 250mL clarified supernatants and mixed thoroughly. This solution was incubated at 4°C overnight, then transferred into sterile 50mL tubes and centrifugated at 4000 RCF for 30 minutes @ 4°C. Supernatants were discarded and pellets resuspended in a total of 1mL 150mM NaCl, 50mM Tris buffer. For final purification steps, cell pellets were combined with PEG-precipitated virus, treated with Benzonase nuclease (250U/uL) for 30 minutes at 37°C, then centrifugated at 7188 RCF for 30 minutes at 4°C. The clarified lysate was transferred to Optiseal tubes (Beckman #361625) containing layered gradients of iodixanol (15%, 25%, 40%, and 54% iodixanol solutions). Tubes were ultracentrifugated in a Beckman 70 Ti fixed-angle rotor at 70,000 rpm for 1hr 40min at 15°C on a WX 80+ centrifuge (Thermo Scientific). The layers containing virus were carefully removed from between the 40% and 54% iodixanol layers, filtered through 0.22um polyethersulfone (PES) membrane syringe filters, and concentrated in Amicon Ultra-15 Centrifuge Units (Millipore #UFC910008), with three full buffer exchanges with DPBS to remove all residual iodixanol. Each batch of virus was concentrated to a final volume of 250-300 µL total, aliquoted in 5 µl aliquots and stored at −80°C, and titered by quantitative PCR.

### Production of rabies viruses

EnvA-enveloped ΔG rabies viruses were made as described previously^30, 88, 89^, using genome plasmids pRVΔG-4Flpo (Addgene 122050) and pRVΔG-4Cre (Addgene 98034).

EnvA-enveloped ΔGL rabies viruses were rescued in 15 cm plates as described^23^ using genome plasmids pRVΔGL-4Flpo (Addgene 98040) and pRVΔGL-4Cre (Addgene 98039). Supernatants from rescue plates were passaged on 15 cm plates of HEK 293T/17 cells (ATCC) transfected with pLV-TTBG and pLV-TTBL (described above) using Xfect transfection reagent (Takara 631318) according to the manufacturer’s protocol. Supernatants were titered on reporter cells^23^ and used to infect BHK-EnvA2-TTBL2 cells (described above) at a multiplicity of infection of approximately two. Supernatants from these producer cells were collected and purified as described^88, 89^.

### Titering of rabies viruses

Recombinase-expressing rabies viruses were titered primarily on the reporter cell lines 293T-FLEX-BC, 293T-F14F15S-BC, 293T-FLEX-BC-TVA and 293T-F14F15S-BC-TVA in infectious units per ml (iu/ml) as described^30^ but using three-fold serial dilutions, instead of ten-fold dilutions, for higher precision. For the corticostriatal experiment, in order to directly compare viral titers in a way that did not depend on the efficacy of the encoded recombinases, we made the BHK-B19L-TVA950 cell line (described above), which expresses both TVA, to allow infection by EnvA-enveloped viruses, and L, in order to allow intracellular replication of ΔGL viruses so that infected cells could be clearly labeled using immunostaining against the rabies virus nucleoprotein. BHK-B19L-TVA950 cells, as well as BHK-B19L cells, were infected with three-fold serial dilutions of RVΔGL-Cre(EnvA) and RVΔGL-Flpo(EnvA. Three days after infection, cells were fixed with 4% paraformaldehyde in PBS and immunostained with a FITC-conjugated anti-nucleocapsid monoclonal antibody blend (Light Diagnostics Rabies DFA, EMD Millipore, catalog # 5100) diluted 1:100 in 1% BSA, 0.1% Triton in PBS. Immunolabeled cells were analyzed by flow cytometry to determine titers as described^88^.

The titers of the rabies viruses were determined to be as follows: RVΔGL-Cre(EnvA):

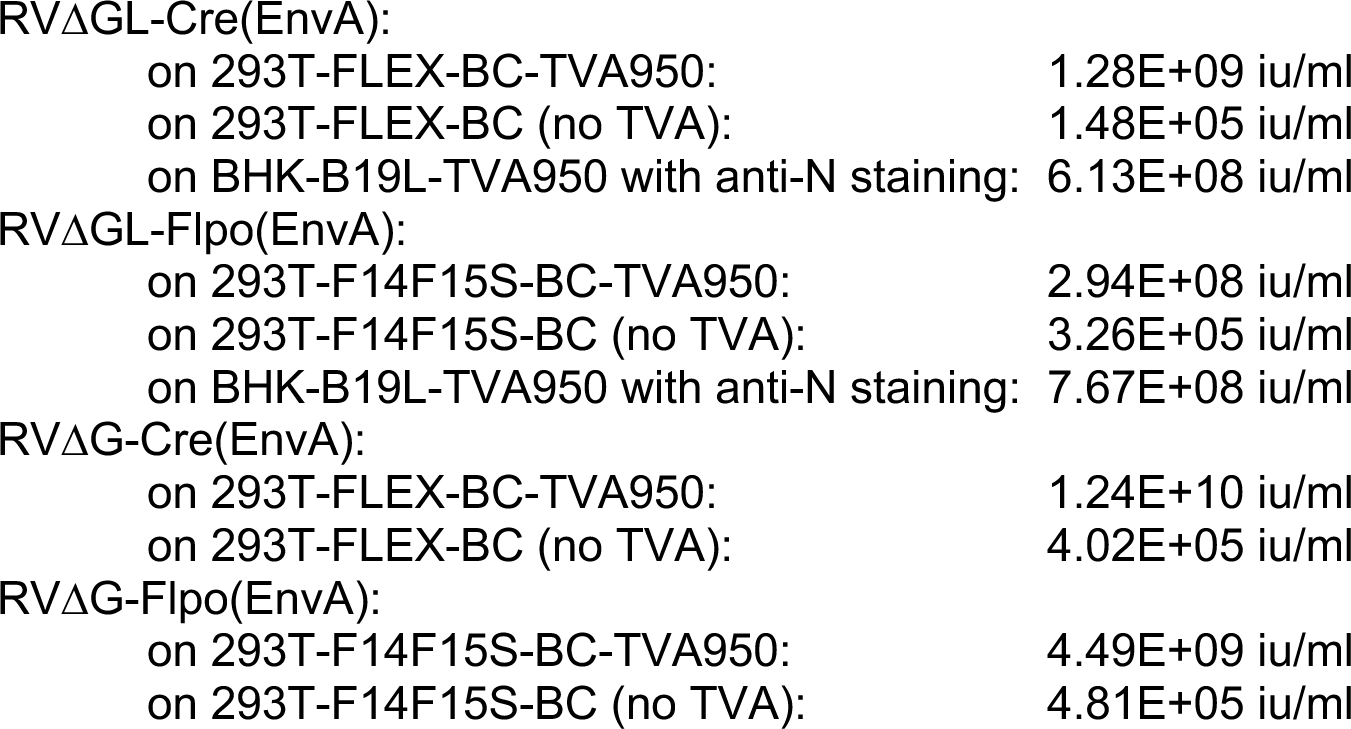
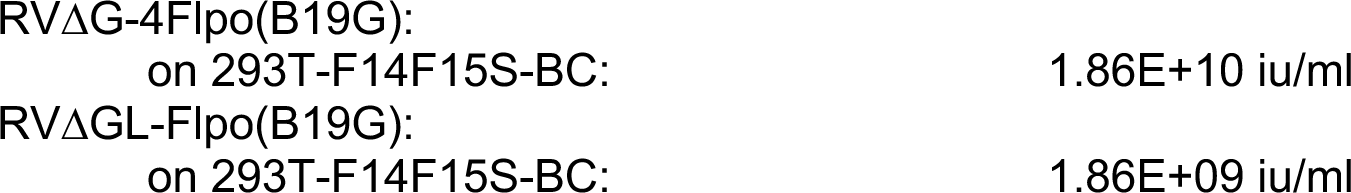

### Stereotaxic injections

Stereotaxic injections were made in anesthetized adult mice of both sexes using a stereotaxic instrument (Stoelting Co., 51925) and a custom injection apparatus consisting of a hydraulic manipulator (Narishige, MO-10) with headstage coupled via custom adaptors to a wire plunger advanced through pulled glass capillaries (Drummond, Wiretrol II) back-filled with mineral oil and front-filled with viral vector solution. We have described this injection system in detail previously^48^.

For the corticostriatal experiments (Fig. 1), 200 nl of AAV2-retro-syn-Flpo (1.16E+13 GC/mL; Ai14 or Ai14 x TRE-CB mice) or AAV2-retro-synP-mCre (1.48E+13 GC/mL, diluted to 1.16E+13 GC/mL for matching to the Flpo version; Ai65F or Ai65F x TRE-CB mice) was injected into dorsolateral striatum (AP = +0.74 mm w.r.t. bregma, LM = 2.25 mm w.r.t. bregma, DV = −2.30 mm w.r.t the brain surface); in the same surgery, 250nl of helper virus mixture (AAV1-syn-F14F15S-sTpEptTA_v2 (diluted to 7.19E+10 GC/mL; Ai14 or Ai14 x TRE-CB mice) or AAV1-syn-FLEX-splitTVA-EGFP-tTA (diluted to 7.22E+10 GC/mL; Ai65F or Ai65F x TRE-CB mice) mixed with AAV1-TREtight-mTagBFP2-B19G (diluted to 6.50E+11) in a 50/50 ratio by volume) was injected into S1BF, layer 5 DLS projection region (AP = −1.55 mm w.r.t. bregma, LM = 3.00 mm w.r.t. bregma, DV = −0.75 mm w.r.t the brain surface). 7 days after AAV injection, 250nl of RVΔGL-Cre(EnvA) (6.13E+08 iu/ml as titered by nucleoprotein staining on L-expressing cells; see above) or RVΔGL-Flpo(EnvA) (7.67E+08 iu/ml, diluted to 6.13E+08 iu/ml) or RVΔG-Flpo(EnvA) (4.49E+09 iu/ml) or RVΔG-Cre(EnvA) (1.24E+10 iu/ml, diluted to 4.49E+09 iu/ml) was injected at the same cortical injection site as the helper viruses. For no-G controls, DPBS was included instead of AAV1-TREtight-mTagBFP2-B19G.

For the anatomical experiments in Cre lines (Fig. 2), 300 nl of helper AAV mixture (AAV1-syn-FLEX-splitTVA-EGFP-tTA (diluted to 7.22E+10 GC/mL) mixed with AAV1-TREtight-mTagBFP2-B19G (diluted to 6.50E+11 GC/mL) in a 50/50 ratio by volume; note that these are the same dilutions as we have recommended previously^48, 49^ but retitered by qPCR in our own lab) was injected into either primary somatosensory cortex or substantia nigra pars compacta (see below for stereotaxic coordinates). Injection coordinates for substantia nigra pars compacta were: anteroposterior (AP) = −3.00 mm with respect to (w.r.t.) bregma, lateromedial (LM) = +1.50 mm w.r.t bregma, dorsoventral (DV) = −4.20 mm w.r.t the brain surface); injection coordinates for somatosensory cortex were: AP = −0.58 mm w.r.t. bregma, LM = 3.00 mm w.r.t. bregma, DV = −1.00 mm w.r.t the brain surface. Seven days after AAV injection, 500 nl (PV-Cre and DAT-IRES-Cre) of RVΔGL-Flpo(EnvA) (7.67E+08 iu/mL) was injected at the same injection site as the AAVs. The experiments in DAT-P2A-Flpo mice (Fig. 3) were done similarly but using the Flp-dependent AAV-syn-F14F15S-EGFP-tTA_v2 (diluted to 7.19E+10 GC/mL) and RVΔG-Cre(EnvA) (1.24E+10 iu/ml, diluted to 4.49E+09 iu/ml).

For longitudinal two-photon imaging for comparison of RVΔG-Flpo(B19G) and RVΔGL-Flpo(B19G), 200 nL RVΔG-Flpo(B19G) (1.86E+10 iu/ml) or RVΔGL-Flpo(B19G) (1.86E+09 iu/ml) was injected into V1 (AP = −2.70 mm w.r.t. bregma, LM = 2.50 mm w.r.t. bregma, DV = −0.26 mm w.r.t. brain surface) into Ai65F mice. Glass windows composed of a 3mm-diameter glass coverslip (Warner Instruments CS-3R) glued (Optical Adhesive 61, Norland Products) to a 5mm-diameter glass coverslip (Warner Instruments CS-5R) were then affixed over the craniotomy with Metabond (Parkell), and custom stainless steel headplates (eMachineShop) were affixed to the skulls around the windows.

For longitudinal two-photon structural imaging of live monosynaptic tracing at the injection site, a 3 mm craniotomy was opened over primary visual cortex (V1). 300 nl of AAV1-syn-FLEX-splitTVA-EGFP-tTA (diluted to 7.22E+10 GC/mL; Ai65F or Ai65F x TRE-CB mice) mixed with AAV1-TREtight-mTagBFP2-B19G (diluted to 6.50E+11) in a 50/50 ratio by volume was injected into V1 (AP = −2.70 mm w.r.t bregma, LM = 2.50 mm w.r.t. bregma, DV = −0.26 mm w.r.t. brain surface), followed by implantation of windows as described above. Seven days after injection of helper AAVs, the coverslips were removed and 100 nl of RVΔGL-Flpo (EnvA) (7.67E+08 iu/mL) was injected at the same site. Coverslips were reapplied and custom stainless steel headplates (eMachineShop) were affixed to the skulls around the windows.

For functional imaging experiments, the V1 injection coordinates were AP = −2.45 mm w.r.t bregma, LM = 2.00 mm w.r.t bregma, DV = −0.26 mm w.r.t. brain surface, and the RVΔGL-Flpo(EnvA) (7.67E+08 iu/mL) was mixed in a 50/50 ratio by volume with AAV1-syn-F14F15S-jGCaMP7s (5.44E+12 GC/ml, diluted 1:10 in DPBS) before injection of 200 nl of the mixture 7d following the helper AAV injection.

### Doxycycline administration

‘No-dox’ mice were fed with regular rodent chow throughout, while ‘dox (food)’ mice were switched to chow containing doxycycline 200 mg/kg (Fisher Scientific, 14-727-450) beginning two weeks after RV injection and maintained on doxycycline chow until perfusion, in order to suppress rabies viral polymerase and glycoprotein expression. ‘Dox (inj+food)’ mice, including those used for the structural two-photon imaging experiment, also received intraperitoneal injections of 100 mg/kg doxycycline every 12 hours for three days, beginning two weeks after RV injection.

### *In vivo* two-photon imaging and image analysis

For longitudinal two-photon imaging of live monosynaptic tracing, injection sites were imaged on a Prairie/Bruker Ultima IV In Vivo two-photon microscope driven by a Spectra Physics Mai-Tai Deep See laser with a mode locked Ti:sapphire laser emitting at a wavelength of 1020 nm for tdTomato. Mice were reanesthetized and mounted via their headplates to a custom frame, with ointment applied to protect their eyes and with a handwarmer maintaining body temperature. One field of view was chosen in each mouse in the area of maximal fluorescent labelling. The imaging parameters were as follows: image size 1024 X 1024 pixels (565.1 μm x 565.1 μm), 0.360 Hz frame rate, dwell time 2.0 μs, 1x optical zoom, Z-stack step size 1 μm. Image acquisition was controlled with Prairie View 5.4 software. Laser power exiting the 20x water-immersion objective (Zeiss, W plan-apochromat, NA 1.0) varied between 20 and 65 mW depending on focal plane depth (Pockels cell value was automatically increased from 450 at the top section of each stack to 750 at the bottom section). For the example images of labeled cells, maximum intensity projections (stacks of 150-400 μm) were made with Fiji software. Cell counting was performed with the ImageJ Cell Counter plugin. When doing cell counting, week 1 tdTomato labelled cells were defined as a reference; remaining week 1 cells were the same cells at later time point that align with week 1 reference cells but the not-visible cells at week 1 (the dead cells). Plots of cell counts were made with Origin 7.0 software (OriginLab, Northampton, MA) or Prism 9 (GraphPad Software, San Diego, California).

For functional two-photon imaging of source cells, injection sites in left-hemisphere V1 was chosen as the imaging area. This imaging was performed using the same microscope (5.356-Hz frame rate, 1024 X 128 pixels, 565.1 μm x 565.1 μm, dwell time 0.8 μs, 1x optical zoom, scan angle 45 degree) with the same objective and laser (at 920 nm) as in the structural imaging experiments. Laser power at the objective ranged from 10 to 65 mW. Calcium imaging data were acquired in supragranular layers (100 to 200 μm deep). The animal was awake, head-fixed and the animal body is free to lay into a customized tube with some holes for air circulation. No behavioral training or reward was given. In order to find the same FOVs in subsequent weeks, surface vasculature map was draw on a notebook for each mouse under microscope with light source (Leica M50) and put ‘*’ marker near the location of FOVs in this map for reference when doing the first imaging session. The cell body images from averaged data taken in the first session provided a template for fine alignment. Visual stimuli were generated in Matlab (R2015R version) with custom software based on Psychtoolbox (http://psychtoolbox.org) and shown on the same LCD screen as in the widefield mapping experiments. Each condition consisted of 2 s of a full-field sine-wave grating drifting in one direction, presented at 80% contrast with spatial frequency of 0.04 cycles/ degree, followed by 2 s of uniform mean luminance (gray). All permutations of 12 directions (30° steps) and 5 temporal frequencies (1, 2, 4, 8 and 15 Hz) were shown, in randomized order. The complete set was repeated 10 times, for a total stimulation period of 40 min per FOV per session. Cells were then manually segmented, and single-cell fluorescence traces were extracted by averaging the fluorescence of all pixels masking the soma by ImageJ (versions: 2-0-0-rc-69) software. The mean ΔF/F over the full 2 s of each stimulus condition was used to calculate orientation tuning curves, with background fluorescence (F) in Δ F/F taken as the value of the trace immediately preceding a condition, averaged over all conditions. The raw calcium traces from cells within individual FOVs (not across FOVs, given different imaging conditions across animals and time points) were sorted by mean fluorescence. For ‘tuned’ cells in Figure 4 panel e, the counts are based on all imaged neurons’ individual tuning curves, plotted in MATLAB; any cell showing response to a preferred orientation (including narrowly tuned neurons and broadly tuned neurons) at any temporal frequency (1Hz, 2Hz, 4Hz, 8Hz, or 15Hz) was counted manually as a tuned cell (see Supplementary File S7 for counts of tuned and untuned cells).

### Brain slice electrophysiology

Coronal brain slices containing the S1 were collected from 12 to 32 week old PV-Cre x Ai65F x TRE-CB het/het/het (ΔGL group) or PV-Cre x Ai65F het/het (ΔG and AAV-only groups) for ex vivo patch clamp recordings. Mice were anesthetized with isoflurane, perfused with ice cold cutting solution, and decapitated using scissors. Brains were extracted and immersed in ice-cold (0–4 °C) sucrose cutting solution containing (in mM) 252 sucrose, 26 NaHCO3, 2.5 KCl, 1.25 NaH2PO4, 1 CaCl2, 5 MgCl2 and 10 glucose, which was oxygenated with 95% O2 and 5% CO2. The brains were trimmed and coronal brain slices (300 µm) were sectioned using a vibratome (VT1200, Leica). After sectioning, slices for patch clamp recordings were transferred to a holding chamber containing oxygenated patch clamp recording medium (aCSF) containing (in mM): 126 NaCl, 2.5 KCl, 1.25 NaH2PO4, 1.3 MgCl2, 2.5 CaCl2, 26 NaHCO3, and 10 glucose, where they were maintained at 32 °C for 30 min before decreasing the chamber temperature to ∼20 °C. Slices were transferred one at a time from the holding chamber to a submerged recording chamber mounted on the fixed stage of an Olympus BX51WI fluorescence microscope equipped with differential interference contrast (DIC) illumination. The slices in the recording chamber were continuously perfused at a rate of 2 ml/min with recording aCSF at room temperature and continuously aerated with 95% O2/5% CO2. Whole-cell patch clamp recordings were performed in the fluorescently-labeled PV+ neurons and principal neurons in the S1 and neighboring S2 region. The presynaptic principal neurons to PV cells in the AAV control group were randomly selected based on the size, morphology, and accommodating firing pattern. Glass pipettes with a resistance of 4-8 MΩ were pulled from borosilicate glass (ID 1.2 mm, OD 1.65 mm) on a horizontal puller (Sutter P-97) and filled with an intracellular patch solution containing (in mM): 130 potassium gluconate, 10 HEPES, 10 phosphocreatine Na2, 4 Mg-ATP, 0.4 Na-GTP, 5 KCl, 0.6 EGTA; pH was adjusted to 7.25 with KOH and the solution had a final osmolarity of 293 mOsm. Series resistance was continuously monitored. Cells were discarded when the series resistance changed more than 20%. Data were acquired using a Multiclamp 700B amplifier, a Digidata 1440 A analog/digital interface, and pClamp 10 software (Molecular Devices). Data were analyzed with Clampfit 10 (Molecular Devices). The membrane capacitance (Cm) and Input resistance (Rm) were calculated from a membrane seal test conducted in voltage-clamp mode, in which 100-ms, 5-mV voltage steps were delivered at a frequency of 5 Hz. In order to evaluate the action potential rheobase, a 1-s, positive current was delivered in current-clamp mode at 10pA steps. Rheobase was defined as the minimum current required to depolarize the membrane potential for action potential firing. Finally, to evaluate the relationship between injected current and firing frequency, we delivered a series of 0.5s current pulses in current-clamp at 50pA steps (50-250pA for principal cells, 50-450pA for PV cells). Statistical comparisons were conducted with unpaired Student’s t-test or with a one- or two-way ANOVA followed by a post hoc Dunnett’s test as appropriate (p < 0.05 with a two-tailed analysis was considered significant).

### Perfusions and histology

1 to 12 weeks (depending on experiment; see main text) after injection of rabies virus, mice were transcardially perfused with 4% paraformaldehyde in phosphate-buffered saline. Brains not used for two-photon imaging were postfixed overnight in 4% paraformaldehyde in PBS on a shaker at 4°C and cut into 50 µm coronal (S1 injections) or parasagittal (SNc injections) sections on a vibrating microtome (Leica, VT-1000S). Sections were collected sequentially into 6 tubes containing cryoprotectant, so that each tube contained a sixth of the collected tissue. For coronal sections, 15 rounds of section were collected. For sagittal sections, 12 rounds were collected. Brains from mice used for two-photon imaging were postfixed in 4% paraformaldehyde/30% sucrose in PBS for two days on a shaker at 4°C, then cut into 50 µm sections on a freezing microtome in a plane approximately tangential to the surface of the brain at the imaged location. Sections were immunostained as described^48^ with the following primary antibodies (as applicable) at the following respective dilutions: chicken anti-GFP (Aves Labs GFP-1020) 1:500, guinea pig anti-parvalbumin (Synaptic Systems 195004) 1:1000, sheep anti-tyrosine hydroxylase (Millipore AB1542)) 1:1000, with secondary antibodies donkey anti-chicken Alexa Fluor 488 (Jackson Immuno 703-545-155) 1:200, donkey anti-guinea pig, AlexaFluor 647 conjugated (Jackson Immuno 706-605-148) 1:200, and donkey anti-sheep, AlexaFluor 647 conjugated (Jackson Immuno 713-605-147)) 1:200. Sections were mounted with Prolong Diamond Antifade mounting medium (Thermo Fisher P36970) and imaged on a confocal microscope (Zeiss, LSM 900).

### Cell counts and statistical analysis

Labeled neurons in striatum (in DAT-IRES-Cre mice), contralateral cortex and thalamus (in PV-Cre mice and for the corticostriatal experiments), and at the injection sites were counted either manually or with the automated Cell Counter function in ImageJ, in every sixth 50-µm section on an epifluorescence microscope (Zeiss Imager.Z2). Coronal sections (corticostriatal experiments and PV-Cre) included sections between 1.2mm and −3.3mm relative to Bregma. Sagittal sections (DAT-IRES-Cre) covered sections 3.6mm to 0.0mm relative to Bregma. Cells expressing EGFP, mTagBFP2, or both, along with tdTomato, were counted by adding separate labels to each and then looking for overlapping cells. Groups were compared using single factor ANOVAs. Symbols denoting significance levels on graphs are as follows: *** = p<0.001; ** = 0.001≤p<0.01; * = 0.01≤p<0.05; n.s. = p≥0.05.

## Reporting Summary

Further information on research design is available in the Nature Research Reporting Summary linked to this paper.

## Data Availability

All cell counts and statistical analyses are provided in Supplementary Information. The novel plasmids described in this paper have been deposited with Addgene with the accession numbers given in Methods. The TRE-CB mouse line is available from the Jackson Laboratory (accession number 036974).

## Acknowledgements

We thank Aurora Burds-Connor and Noranne Enzer of the Preclinical Modeling Facility at the Koch Institute for Integrative Cancer Research at MIT for generation of the TRE-CB line from ES cells. We thank Karel Svoboda for advice on titrating jGCaMP6s expression for reduced toxicity. We thank Ayano Matsushima for examination of labeled regions in the DAT-P2A-Flpo mice. Research reported in this publication was supported by BRAIN Initiative awards U01MH106018, U01MH114829, U19MH114830, and RF1MH120017 from the National Institute of Mental Health.

## Author contributions

L.J. contributed molecular biology, cell culture, mouse colony management, virus injections, two-photon imaging, histology, microscopy, data analysis, and preparation of figures. H.A.S. produced viruses and conducted cell culture assays. M.Z. contributed perfusions, histology, microscopy, cell counts, data analysis, and mouse colony management. T.K.L. contributed virus injections, perfusions, histology, microscopy, and mouse colony management. M.M., C.X. and Y.H contributed cloning and molecular biology. N.E.L., M.P., & K.R.B. contributed mouse colony management, assisted with molecular biology. L.R. contributed cell counts. T.L.D. and H.Z. contributed embryonic stem cell targeting for making the TRE-CB mouse line. J.I., M.H., and M.S. assisted with establishing the structural and functional two-photon imaging. X.F and G.F. planned slice electrophysiology experiments; X.F. conducted slice electrophysiology experiments and identified the problem with the original Flp-dependent helper AAV that was used for the original version of the manuscript. I.R.W. designed experiments and wrote the paper with input from L.J., H.A.S., M.Z., and other authors.

## Additional information

Supplementary information is available for this paper.

A preprint version of this paper is available on bioRxiv.

