## Supplementary Figures for "Long-term labeling and imaging of synaptically-connected neuronal networks *in vivo* using double-deletion-mutant rabies viruses"

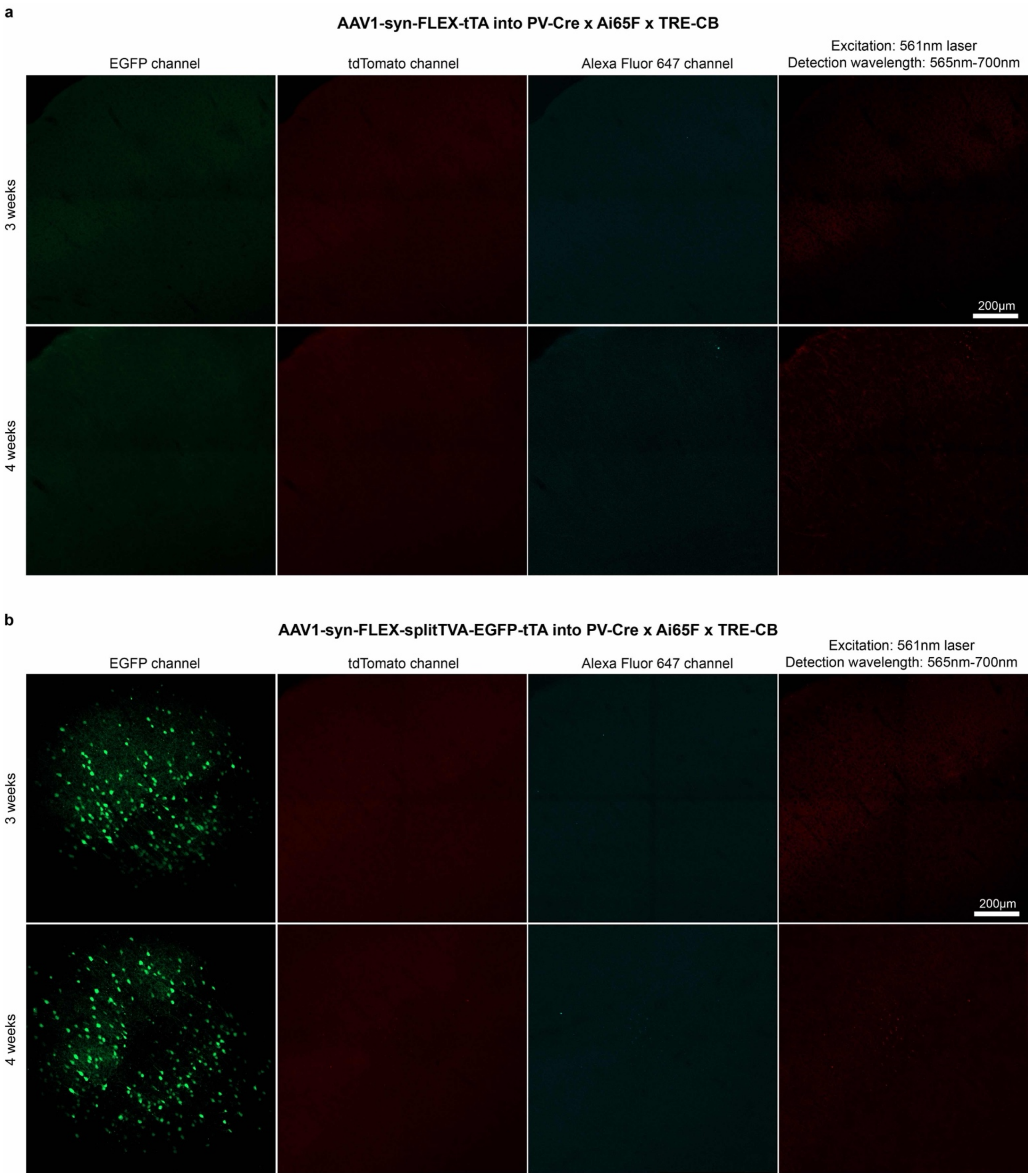


**Extended Data Fig. S1: Intrinsic mCardinal fluorescence was not detected by confocal imaging, suggesting very low L expression in the conditions implemented.** Representative confocal images of injection sites in PV-Cre x Ai65F x TRE-CB mice, showing lack of clear mCardinal fluorescence induced by either of the laser lines (561 nm or 640 nm) flanking the excitation maximum on the Zeiss LSM 900 confocal microscope. Because of the lack of fluorescence in the channel otherwise used for AlexaFluor 647, specifically, we were able to use that channel for immunostaining for PV and TH (Fig. 1 and Extended Data Figs. wasFigS2 & wasFigS3). **a,** Injection sites of a FLEX AAV expressing tTA without a fluorophore, and **b,** injection sites of the FLEX AAV expressing TVA, EGFP, and tTA, with survival times of 3 and 4 weeks in each case. Left: EGFP channel; center-left: tdTomato channel; center-right: AlexaFluor 647 channel; right: "best case" for imaging mCardinal: excitation with 561 nm and collecting all emitted light between 565 nm and 700 nm. Scale bars: 200 μm, apply to all panels.

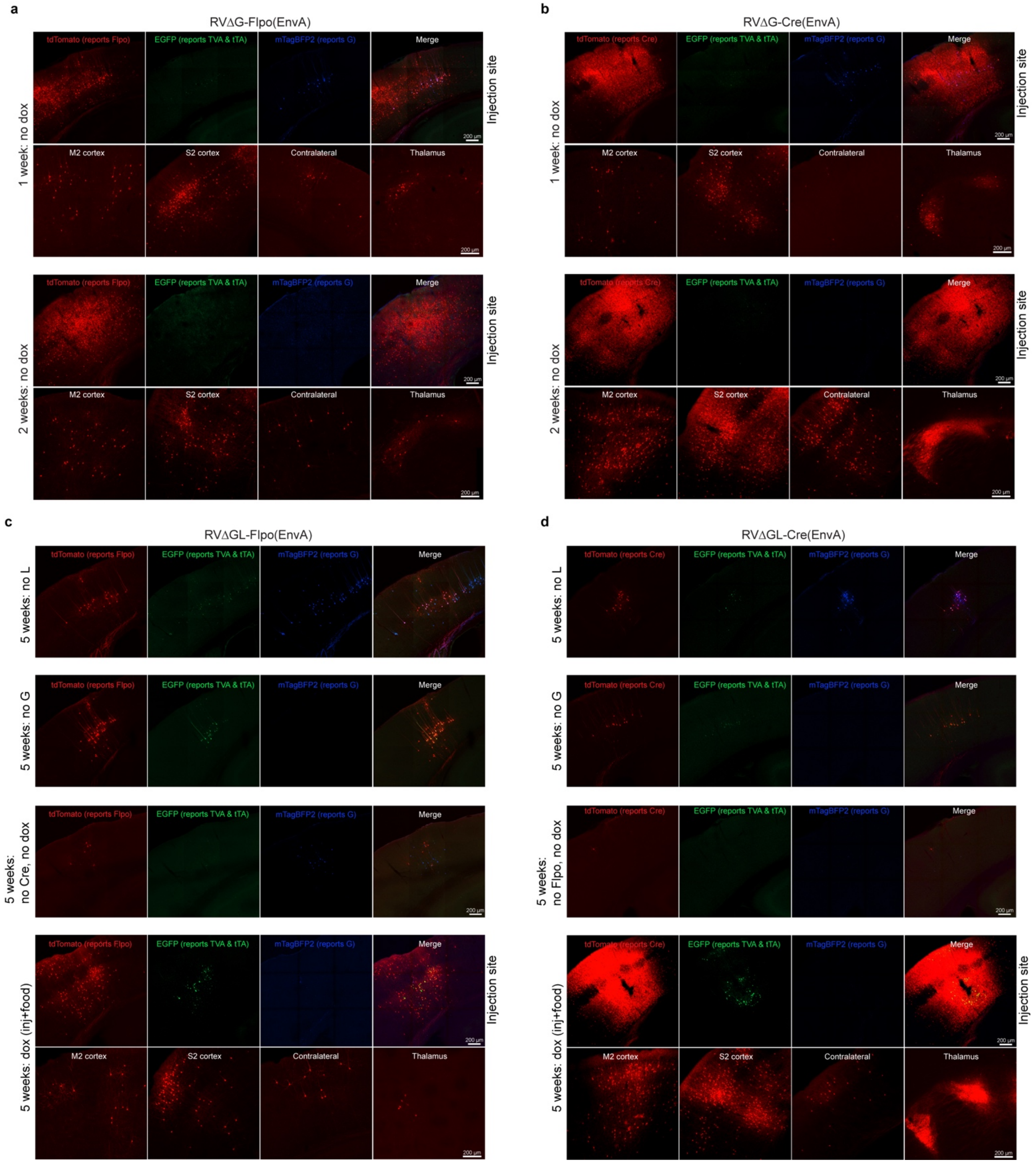


**Extended Data Fig. S2: Representative images of control experiments for corticostriatal experiments.** See Fig. 2 for explanation of experimental design and quantification. **a-b,** First-generation controls (7d and 14d survival times). **c-d,** Controls using second-generation viruses: no L, no G, and no Cre/Flpo controls (all without doxycycline), as well as the 'dox (inj+food)' condition for comparison. Scale bars: 200 μm, apply to all images.

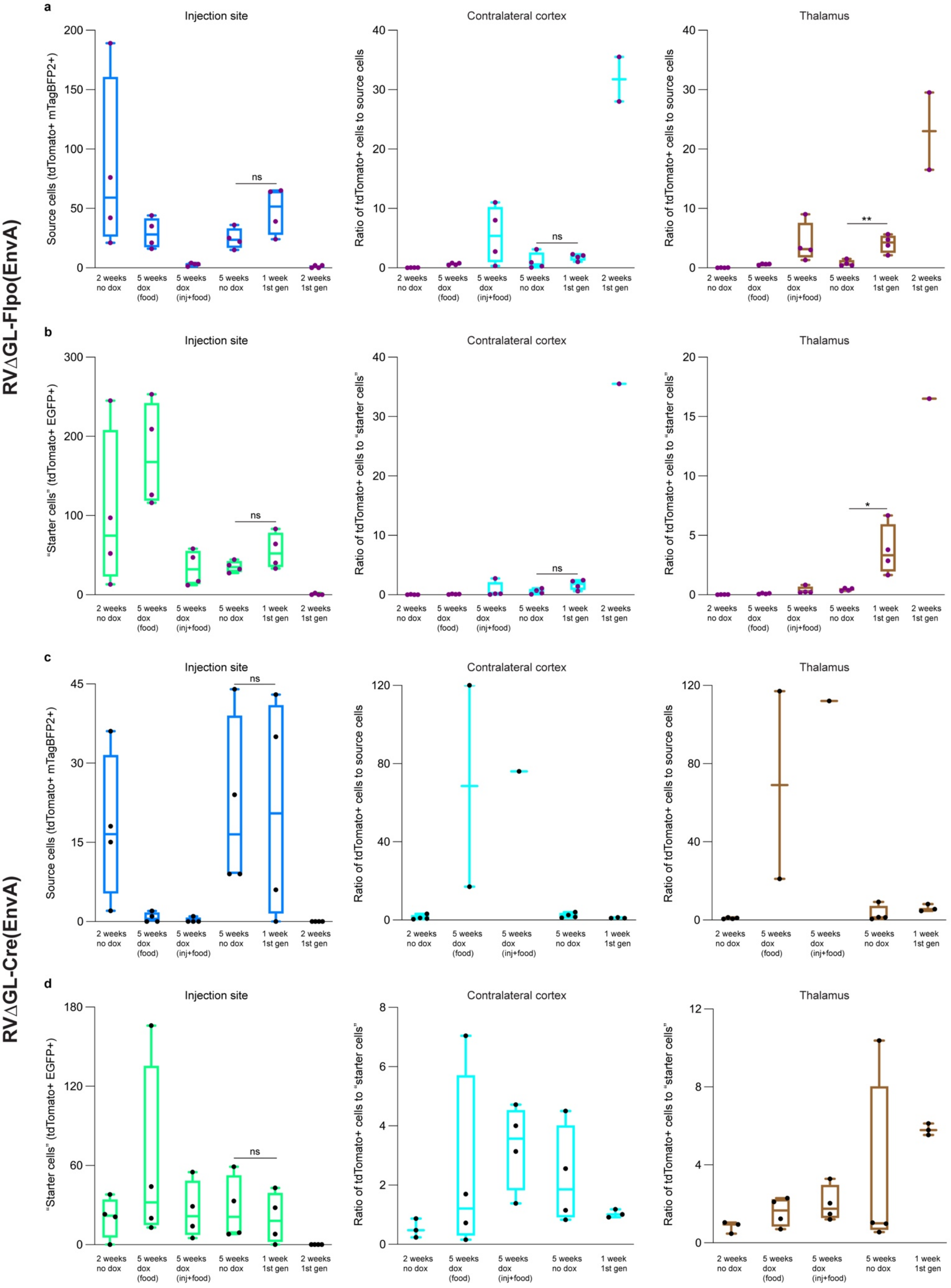


**Extended Data Fig. S3: Counts of corticothalamic "source" and "starter" cells and ratios of contralateral and thalamic cells to source and starter cells.** Counts are totals found in every sixth 50-μm section spanning each brain. **a-b,** Results using RVΔGL-Flpo(EnvA). **a,** Counts of source cells (left), defined as cells coexpressing tdTomato (marking activity of Flpo) and mTagBFP2 (marking expression of G), and ratios of these numbers of source cells to numbers of tdTomato+ cells in contralateral cortex (middle) and thalamus (right). Note the near-absence of source cells in the first-generation conditions at two weeks, suggesting that most had died by this time point. Note also that the low numbers of source cells in the second-generation conditions with doxycycline could reflect either cell death or simply the intended suppression of mTagBFP2 expression in these mice. For mice in which no source cells were found (either because they had died or because mTagBFP2 was suppressed by doxycycline), no ratio is included on the graphs. **b,** Counts and ratios of starting cells, defined here as cells coexpressing tdTomato (marking activity of Flpo) and EGFP (marking expression of TVA and tTA). **c-d,** Results using RVΔGL-Cre(EnvA). **c,** Counts and ratios of source cells, defined here as cells coexpressing tdTomato (marking activity of Cre) and mTagBFP2 (marking expression of G). **d,** Counts and ratios of starting cells, defined here as cells coexpressing tdTomato (marking activity of Cre) and EGFP (marking expression of TVA and tTA).

*(images in external file)*

**Extended Data Fig. S4: Series of whole-brain images of labeled inputs to corticostriatal neurons using first- and second-generation rabies viral vectors expressing Cre.** Images are tiled confocal images of every sixth 50-μm coronal section spanning most of the rostrocaudal extents of the brains.

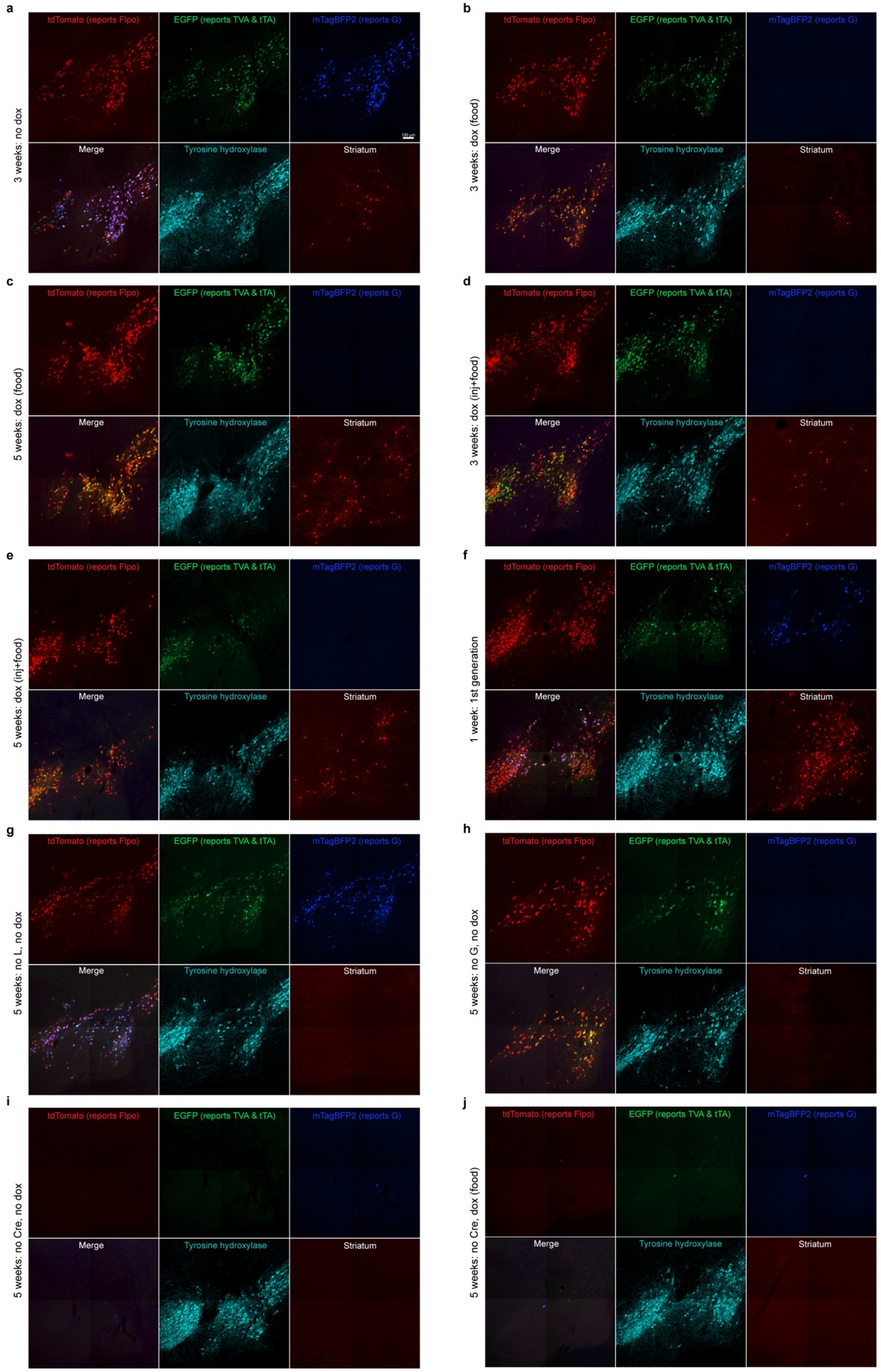
a
3 weeks: no dox
b
3 weeks: dox (food)
c
5 weeks: dox (food)
d
3 weeks: dox (inj+food)
e
5 weeks: dox (inj+food)
f
1 week: 1st generation
g
5 weeks: no L, no dox
h
5 weeks: no G, no dox
i
5 weeks: no Cre, no dox
j
5 weeks: no Cre, dox (food)
tdTomato (reports Flpo)
EGFP (reports TVA & tTA)
mTagBFP2 (reports G)
Merge
Tyrosine hydroxylase
Striatum
100 µm

**Extended Data Fig. S5: Representative images of second-generation monosynaptic tracing of inputs to dopaminergic midbrain neurons using RVΔGL-Flpo.** See Fig. 1 for explanation of experimental design and for quantification. **a,** 3 weeks, no doxycycline, **b,** 3 weeks, with doxycycline, **c,** 5 weeks, with doxycycline. Control conditions shown here: **d,** first-generation (ΔG) system, **e,** no L, **f,** no G, **g,** no Cre, no doxycycline, **h,** no Cre, with doxycycline. Note that omitting L gave very similar results to omitting G.

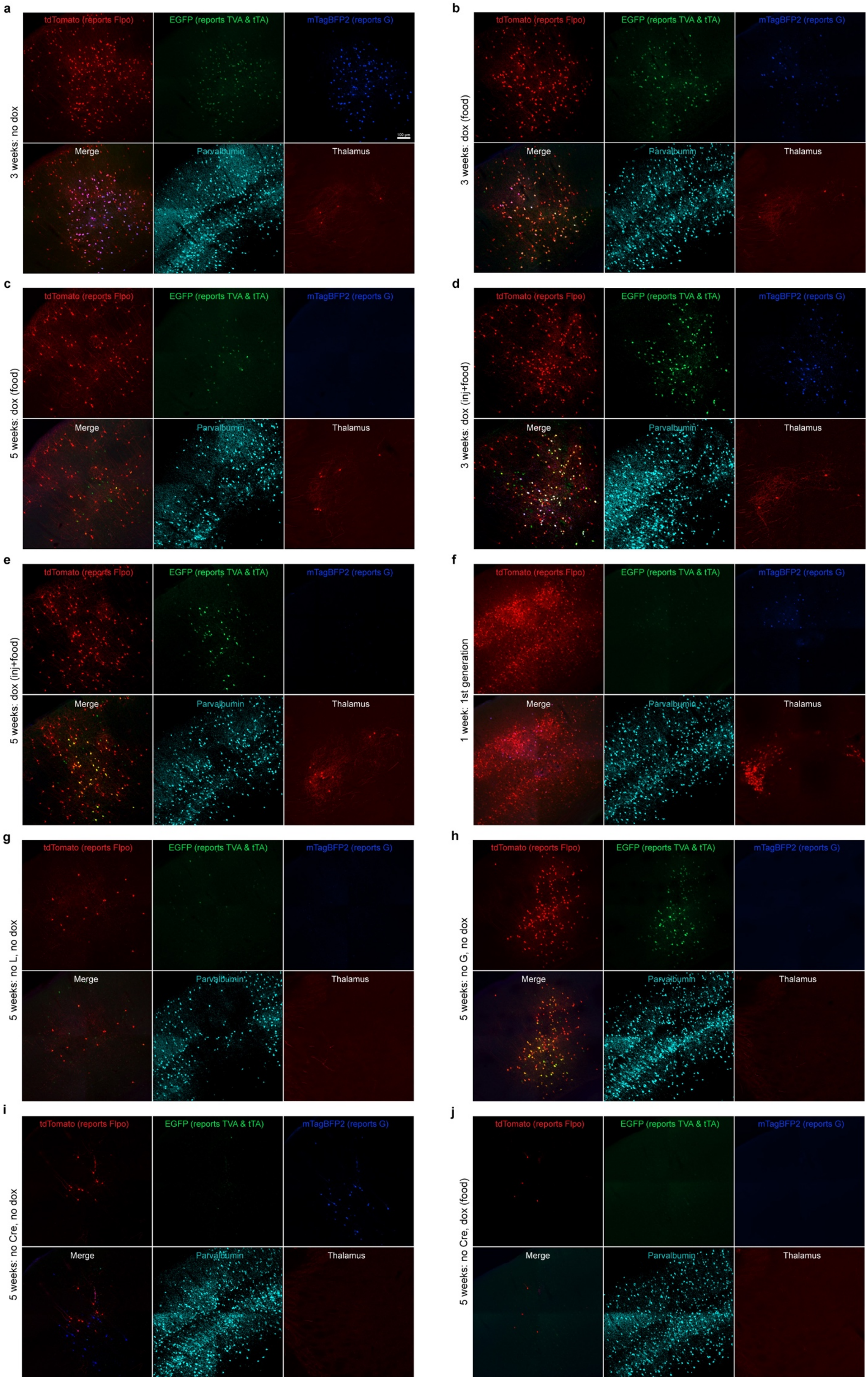
a
3 weeks: no dox
b
3 weeks: dox (food)
c
5 weeks: dox (food)
d
3 weeks: dox (inj+food)
e
5 weeks: dox (inj+food)
f
1 week: 1st generation
g
5 weeks: no L, no dox
h
5 weeks: no G, no dox
i
5 weeks: no Cre, no dox
j
5 weeks: no Cre, dox (food)
tdTomato (reports Flpo)
EGFP (reports TVA & tTA)
mTagBFP2 (reports G)
Merge
Parvalbumin
Thalamus

**Extended Data Fig. S6: Representative images of second-generation monosynaptic tracing of inputs to parvalbumin-expressing cortical interneurons.** See Fig. 1 for explanation of experimental design and for quantification. **a,** 3 weeks, no doxycycline, **b,** 3 weeks, with doxycycline (note that there is still blue fluorescence in this image, suggesting that, at one week after the mice were switched to dox food, the doxycycline has not completely suppressed G and presumably L expression), **c,** 5 weeks, with doxycycline. Control conditions shown here: d) first-generation (ΔG) system, e) no L, f) no G, g) no Cre, no doxycycline, h) no Cre, with doxycycline. Again, omitting L and omitting G gave very similar results.

*(images in external file)*

**Extended Data Fig. S7: Whole-hemisphere image series of labeled inputs to dopaminergic neurons using first- and second-generation rabies viral vectors expressing Cre.** Images are tiled confocal images of every sixth 50-μm parasagittal section spanning the injected hemispheres.

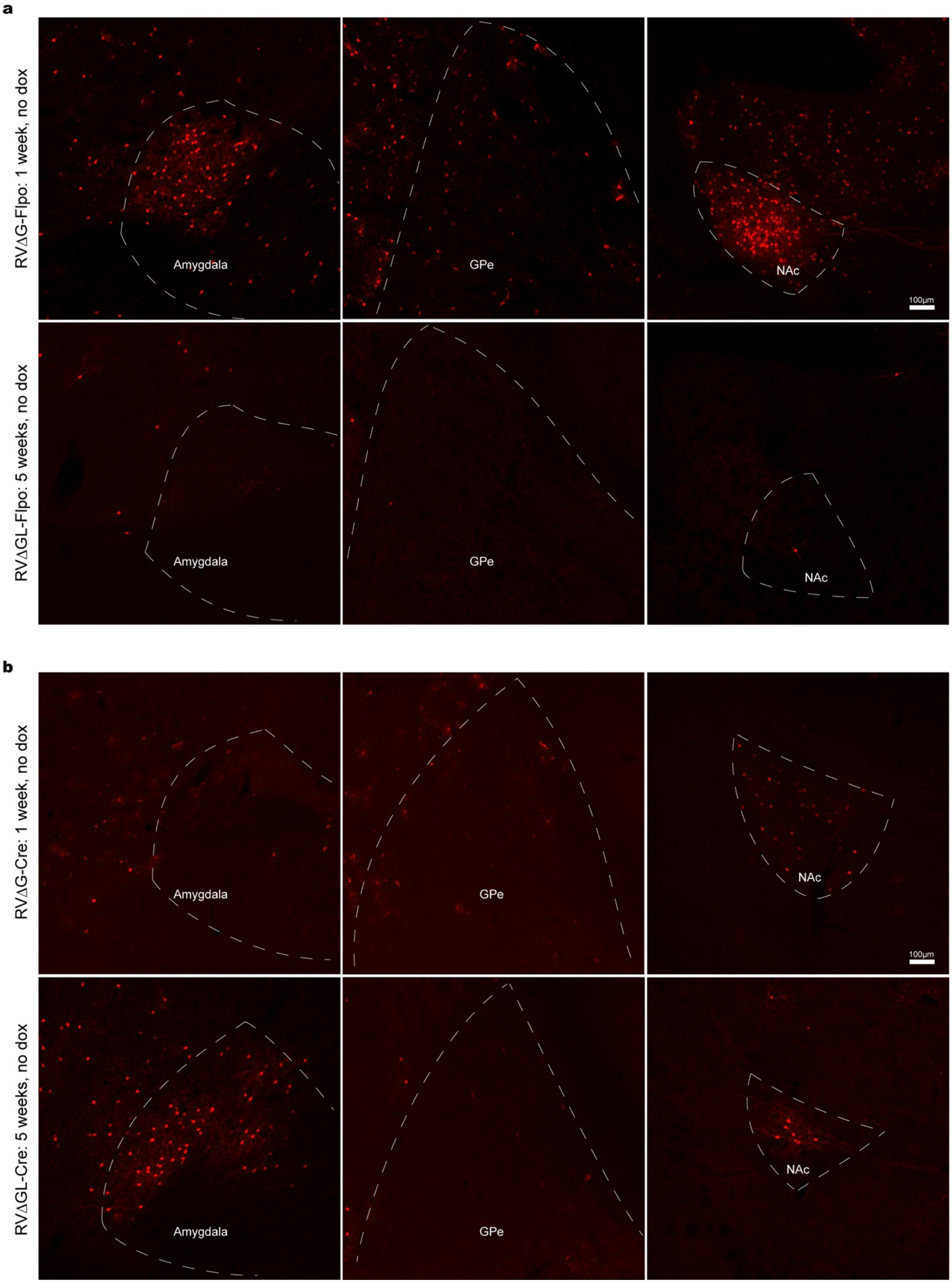


**Extended Data Fig. S8: Examples of extrinsic inputs to dopaminergic cells labeled with first- and second-generation RV vectors expressing Flpo and Cre.**

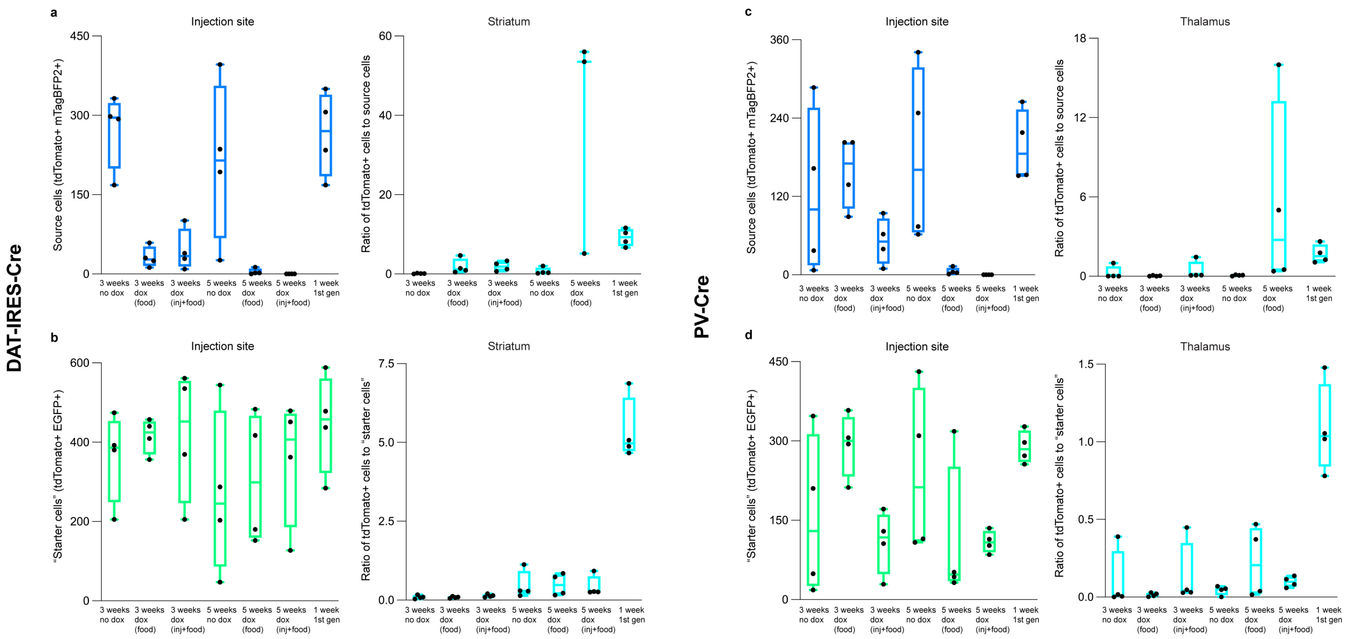


**Extended Data Fig. S9: Counts of dopaminergic and parvabuminergic source cells and ratios of striatal and thalamic cells to starting cells in Cre mice.** Counts are totals found in every sixth 50-μm section spanning each brain. **a-b,** Results using RVΔGL-Flpo(EnvA) in DAT-IRES-Cre. **a,** Counts of source cells (left), defined as cells coexpressing tdTomato (marking activity of Flpo) and mTagBFP2 (marking expression of G), and ratios of these numbers of source cells to numbers of tdTomato+ cells in striatum (right). Note that the low numbers of source cells in the second-generation conditions with doxycycline could reflect either cell death or simply the intended suppression of mTagBFP2 expression in these mice. Note also that there is no graph for the ratio of labeled striatal cells to source cells for the "5 weeks dox (inj + food)" because no source cells were found for that condition (i.e., doxycycline appears to have fully suppressed mTagBFP2 expression). **b,** Counts and ratios of starter cells, defined here as cells coexpressing tdTomato (marking activity of Flpo) and EGFP (marking expression of TVA and tTA). **c-d,** Results using RVΔGL-Flpo(EnvA) in PV-Cre. **c,** Counts and ratios of starting cells, defined here as cells coexpressing tdTomato (marking activity of Flpo) and mTagBFP2 (marking expression of G). Note that again there is no graph for the ratio of labeled contralateral cells to source cells for the "5 weeks dox (inj + food)" because no source cells were found for that condition. **d,** Counts and ratios of starting cells, defined here as cells coexpressing tdTomato (marking activity of Flpo) and EGFP (marking expression of TVA and tTA).

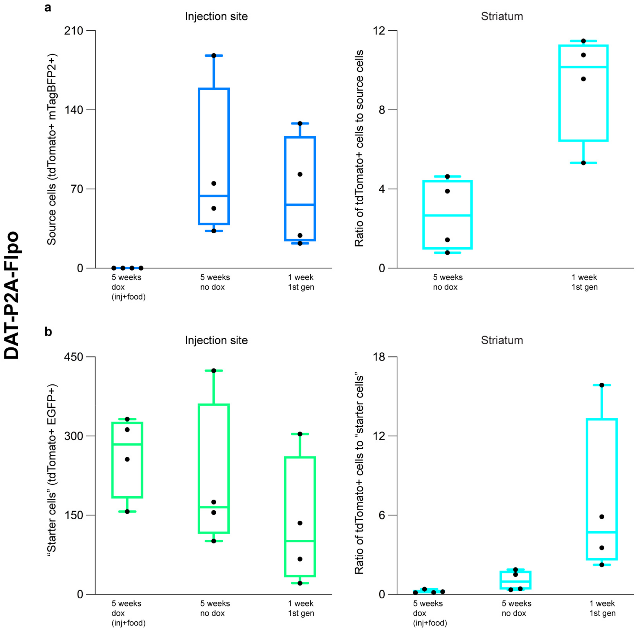


**Extended Data Fig. S10: Counts of dopaminergic and parvabuminergic source cells and ratios of striatal and thalamic cells to starting cells using RVΔGL-Cre(EnvA) in DAT-P2A-Flpo mice.** Counts are totals found in every sixth 50-μm section spanning each brain. **a,** Counts of source cells (left), defined as cells coexpressing tdTomato (marking activity of Cre) and mTagBFP2 (marking expression of G), and ratios of these numbers of source cells to numbers of tdTomato+ cells in striatum (right). Note that the low numbers of source cells in the second-generation condition with doxycycline could reflect either cell death or simply the intended suppression of mTagBFP2 expression in these mice. **b,** Counts and ratios of starter cells, defined here as cells coexpressing tdTomato (marking activity of Cre) and EGFP (marking expression of TVA and tTA).

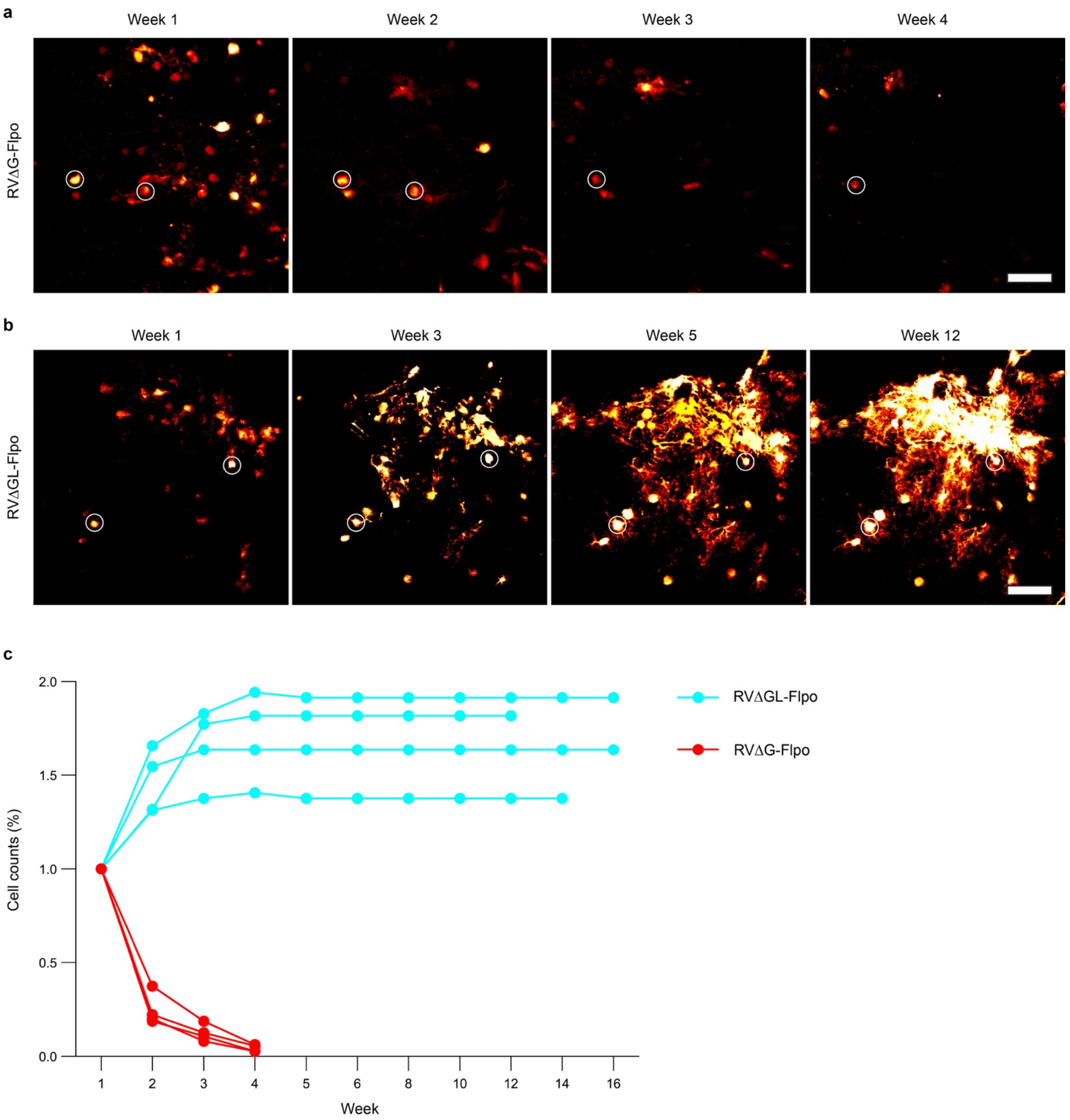


**Extended Data Fig. S11: Second-generation rabies virus encoding Flpo does not kill most labeled neurons for at least 16 weeks. a,** Representative images of longitudinal two-photon structural imaging fields of view (FOVs) from one week (left) to 4 weeks (right) after injection of a first-generation rabies viral vector encoding Flpo (RVΔG-Flpo). The viruses in this experiment were coated with the native RV glycoprotein for direct (TVA-independent) infection of neurons. Images are of the same FOV at different time points. This first-generation virus killed almost all infected cells in this FOV within 4 weeks: only two labeled cells (circled) survived until the 4-week timepoint. Scale bar: 100 μm. **b,** Representative images of longitudinal two-photon structural imaging FOVs from one week (left) to 12 weeks (right) after injection of a second-generation rabies viral vector encoding Flpo (RVΔGL-Flpo). Images are of the same FOV at different time points. All labeled cells are still present at 16 weeks, the last timepoint of imaging. Two example labeled cells are circled as fiducial markers. Scale bar: 100 μm.

**c,** Fraction of visible labeled cells over time, relative to the number visible at one week after RVΔG-Flpo or RVΔGL-Flpo injection; connected sets of dots represent counts obtained from the same FOV within the same mouse at the different time points (for RVΔG-Flpo: 1 week to 4 weeks; for RVΔGL-Flpo: 1 week to 16 weeks). Cells infected by RVΔG-Flpo (red) have almost entirely disappeared by 4 weeks postinjection. The number of cells labeled by RVΔGL-Flpo (cyan) increases up to 4 weeks postinfection and then remains constant for as long as the brains were imaged.

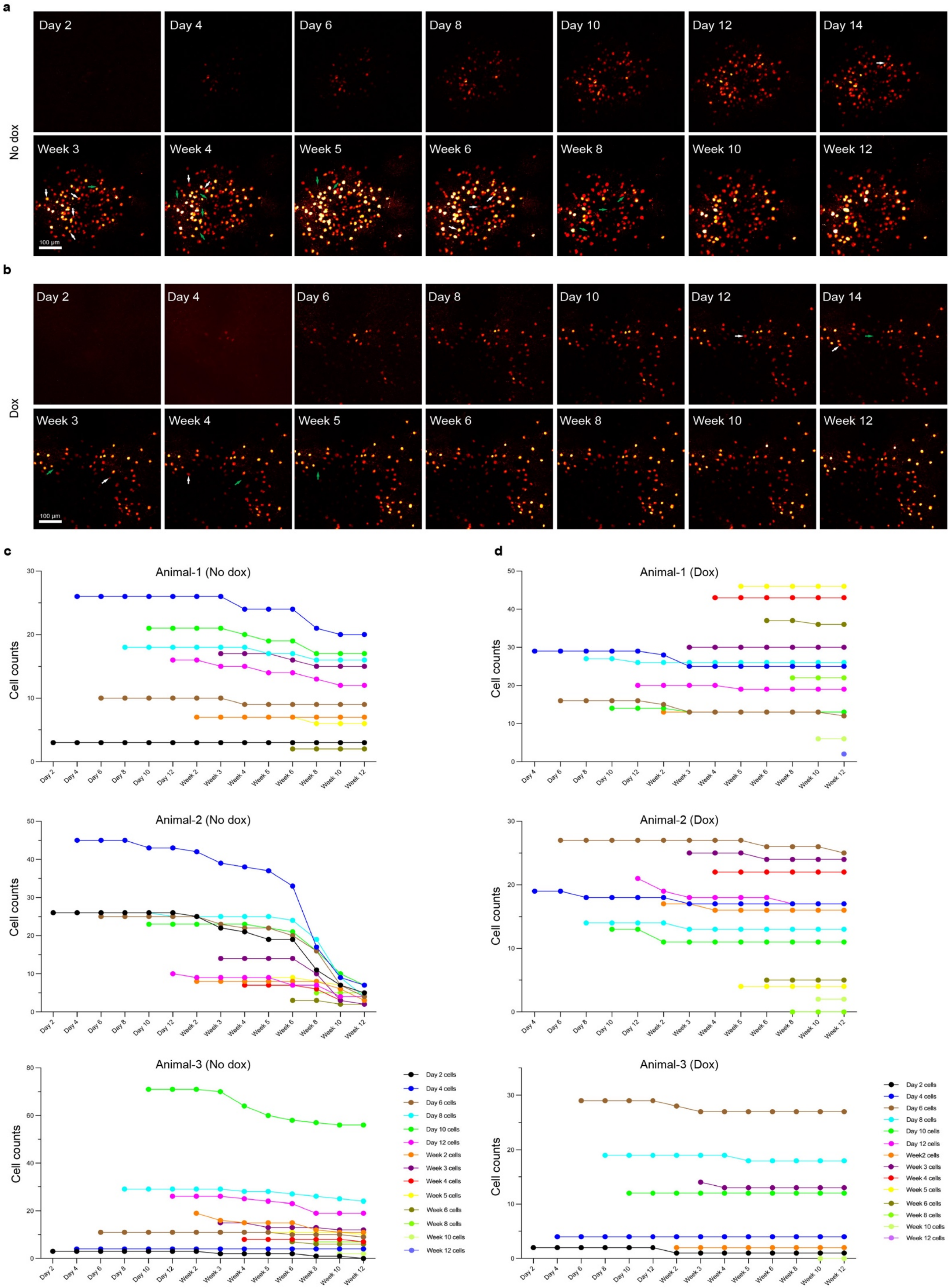
a
No dox
Day 2
Day 4
Day 6
Day 8
Day 10
Day 12
Day 14
Week 3
Week 4
Week 5
Week 6
Week 8
Week 10
Week 12
100 µm
b
Dox
Day 2
Day 4
Day 6
Day 8
Day 10
Day 12
Day 14
Week 3
Week 4
Week 5
Week 6
Week 8
Week 10
Week 12
100 µm
c
Animal-1 (No dox)
Animal-2 (No dox)
Animal-3 (No dox)
d
Animal-1 (Dox)
Animal-2 (Dox)
Animal-3 (Dox)
Cell counts
Day 2
Day 4
Day 6
Day 8
Day 10
Day 12
Week 2
Week 3
Week 4
Week 5
Week 6
Week 8
Week 10
Week 12
Day 2 cells
Day 4 cells
Day 6 cells
Day 8 cells
Day 10 cells
Day 12 cells
Week 2 cells
Week 3 cells
Week 4 cells
Week 5 cells
Week 6 cells
Week 8 cells
Week 10 cells
Week 12 cells

**Extended Data Fig. S12: Additional examples of longitudinal two-photon imaging of second-generation monosynaptic tracing *in vivo*. a-b,** Example images of tdTomato-labelled neurons at the injection site in V1 over all imaging sessions, from 2 days to 12 weeks, in two mice. The ‘No dox’ mice (**a**) were fed with regular food throughout, while the ‘Dox’ mice (**b**) were fed with food containing doxycycline (200 mg/kg) starting at two weeks after RV injection until perfusion at week 12, in order to suppress expression of the rabies virus polymerase and glycoprotein genes. White arrows show example cells that are last seen at that time point, with green arrows indicating the former positions of those now-missing cells at the next time point. In these examples, a total of 20 out of 141 cells were lost in the ‘No dox’ mouse, whereas 5 out of 98 cells were lost in the ‘Dox’ mouse. Scale bar: 100 µm. **c-d,** Counts of tdTomato-labeled cells appearing at each timepoint in individual mice. Each connected set of dots represents the numbers of the cells that appeared at one timepoint that are still present at the subsequent timepoints.

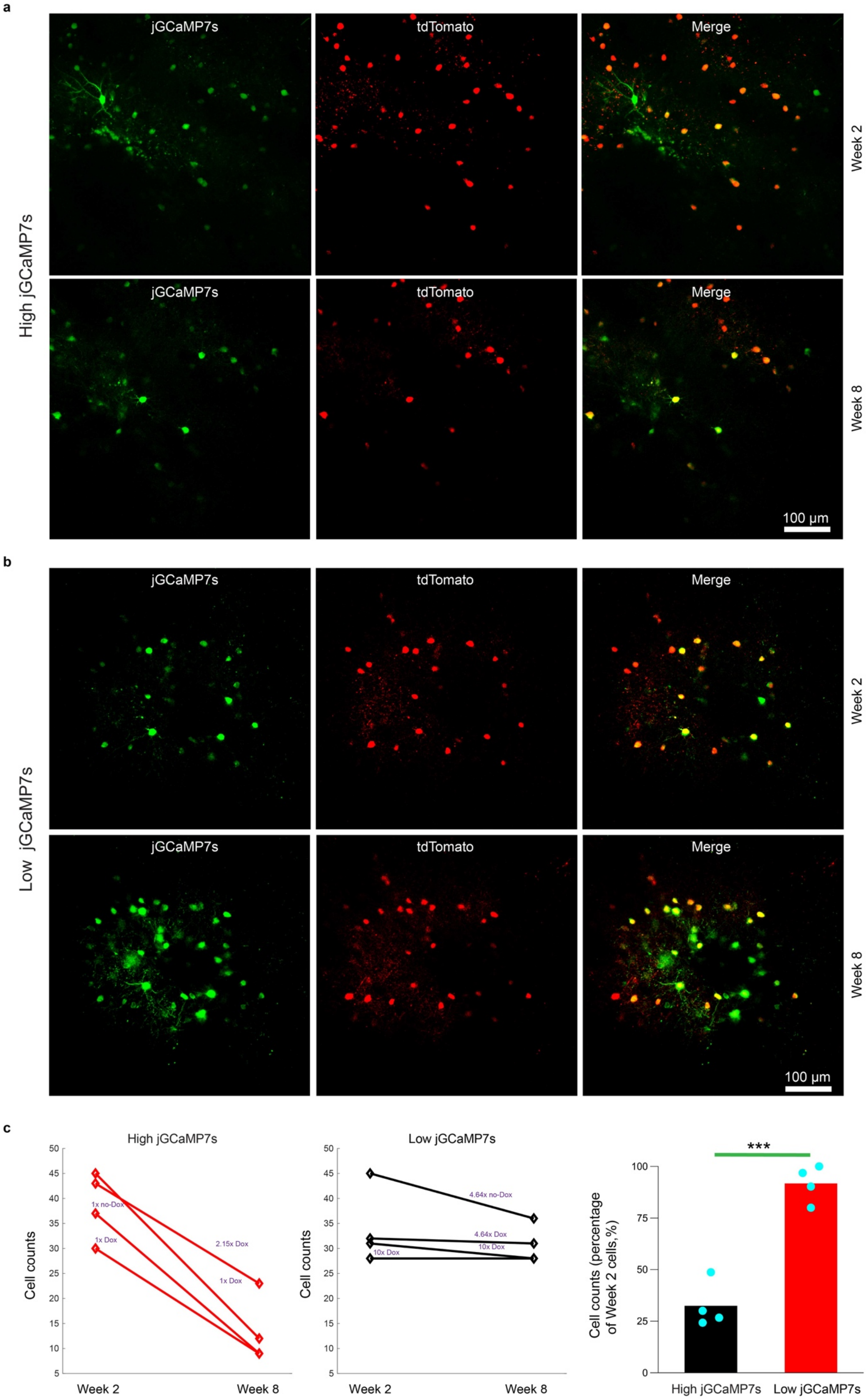
a
jGCaMP7s
tdTomato
Merge
Week 2
Week 8
High jGCaMP7s
100 µm
b
Low jGCaMP7s
c
High jGCaMP7s
Low jGCaMP7s
Cell counts
Week 2
Week 8
1x no-Dox
1x Dox
2.15x Dox
4.64x no-Dox
4.64x Dox
10x Dox
Cell counts (percentage of Week 2 cells,%)
***
High jGCaMP7s
Low jGCaMP7s

**Extended Data Fig. S13: Titration of jGCaMP7s AAV concentration to minimize toxicity from jGCaMP7s overexpression. a,** Representative two-photon images of neurons expressing tdTomato and jGCaMP7s, with the Flp-dependent AAV1 expressing jGCaMP7s injected at a higher concentration (2.15x diluted from maximum), with doxycycline administered. Top row: 2 weeks after injection. Bottom row: the same FOV at 8 weeks after injection. Numerous cells visible at 2 weeks are no longer seen at 8 weeks. Scale bar: 100 um. **b,** Representative two-photon images from another mouse, in which the Flp-dependent AAV1 expressing jGCaMP7s had been injected at a lower concentration (4.64x diluted from maximum), with doxycycline administered. Top row: 2 weeks after injection. Bottom row: the same FOV at 8 weeks after injection. Almost all cells seen at 2 weeks are still present at 8 weeks. Scale bar: 100 µm. **c,** Counts of jGCaMP7s- and tdTomato-expressing neurons infected with high-concentration (left graph) and low-concentration (middle graph) jGCaMP7s AAV. Each data point was obtained from one FOV from each animal. Virus dilution factor and doxycycline condition are marked on the top of each line; for example, ‘10x, Dox’ means the mouse received the 10-fold dilution of jGCaMP7s AAV and doxycycline food two weeks after the second injection. The surviving cell percentage is shown in the right graph; there is a significant decrease in the high concentration of jGCaMP7s group (***$p=0.00036<0.001$, one-way ANOVA test).

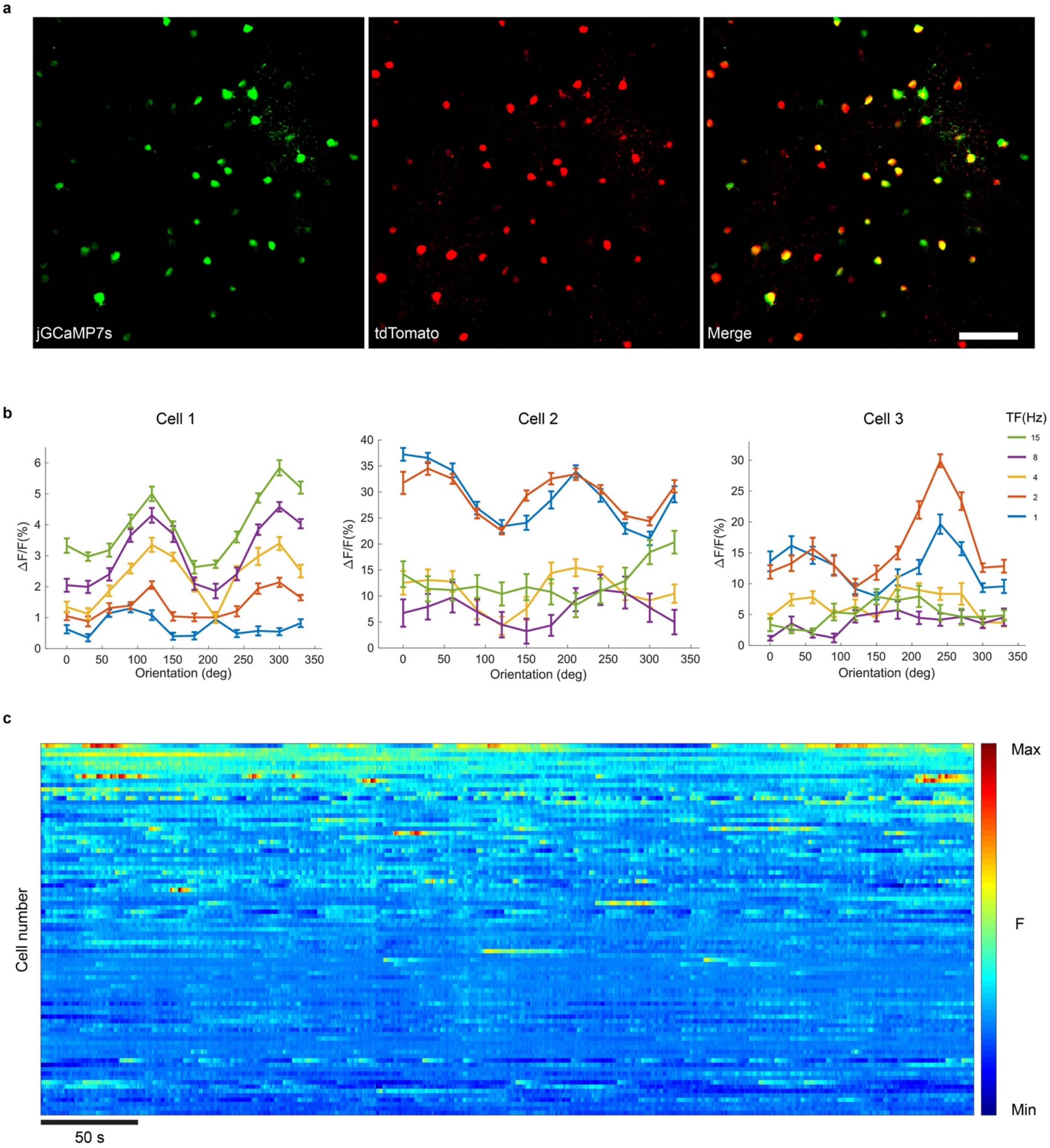


**Extended Data Fig. S14: Additional examples of two-photon functional imaging of labeled cells in mouse visual cortex. a,** Representative two-photon images of neurons expressing jGCaMP7s and tdTomato at 3 weeks. Scale bar: 100 µm. **b,** Direction tuning curves of three different jGCaMP7s-expressing neurons obtained with drifting gratings presented at 12 directions of motion and 5 temporal frequencies (TF), repeated 10 times (mean ΔF/F ± s.e.m.) at 8 weeks; color coding and axis labels are as in Fig. 4. **c,** Single-cell fluorescence time courses for 85 cells over the first 480 s of visual stimulation, showing robust spontaneous and evoked activity. Scale bar: 50 s.

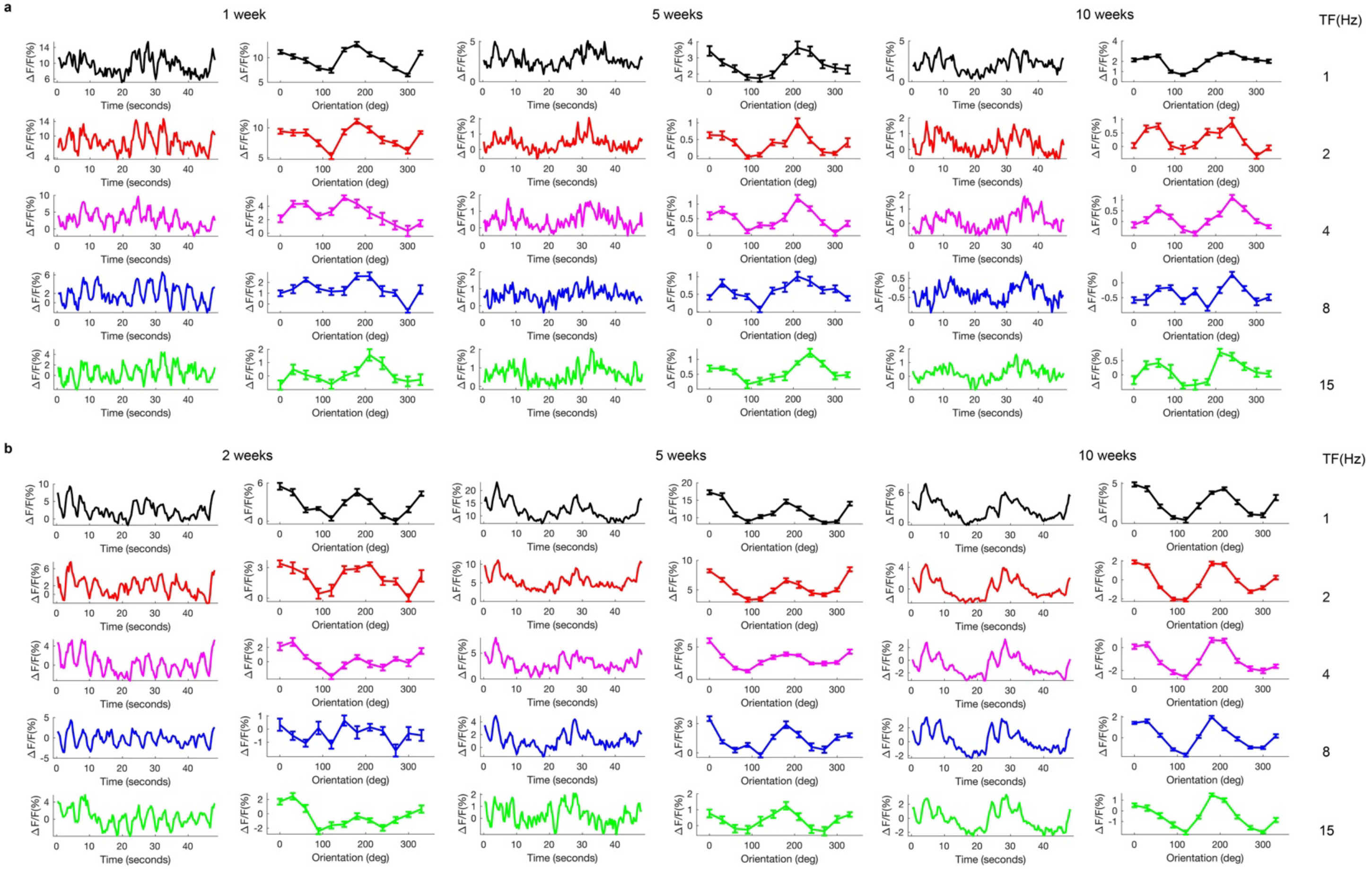


**Extended Data Fig. S15: jGCaMP7s signals and tuning curves of two example V1 neurons over multiple imaging sessions. a,** jGCaMP7s signals and tuning curves of a "week 1" cell (i.e., in which jGCaMP7s and tdTomato fluorescence could be detected seven days after RVΔGL-Flpo(EnvA) and AAV1-syn-F14F15S-jGCaMP7s injection). These data were obtained with drifting gratings presented at 12 directions of motion and 5 temporal frequencies (TF), repeated 10 times (tuning curve: mean ΔF/F ± s.e.m; jGCaMP7s signals: mean ΔF/F) in three different imaging sessions (left: week 1; middle: week 5; right: week 10). **b,** jGCaMP7s signals and tuning curves of a "week 2" cell (the same cell as used for Fig. 4d).

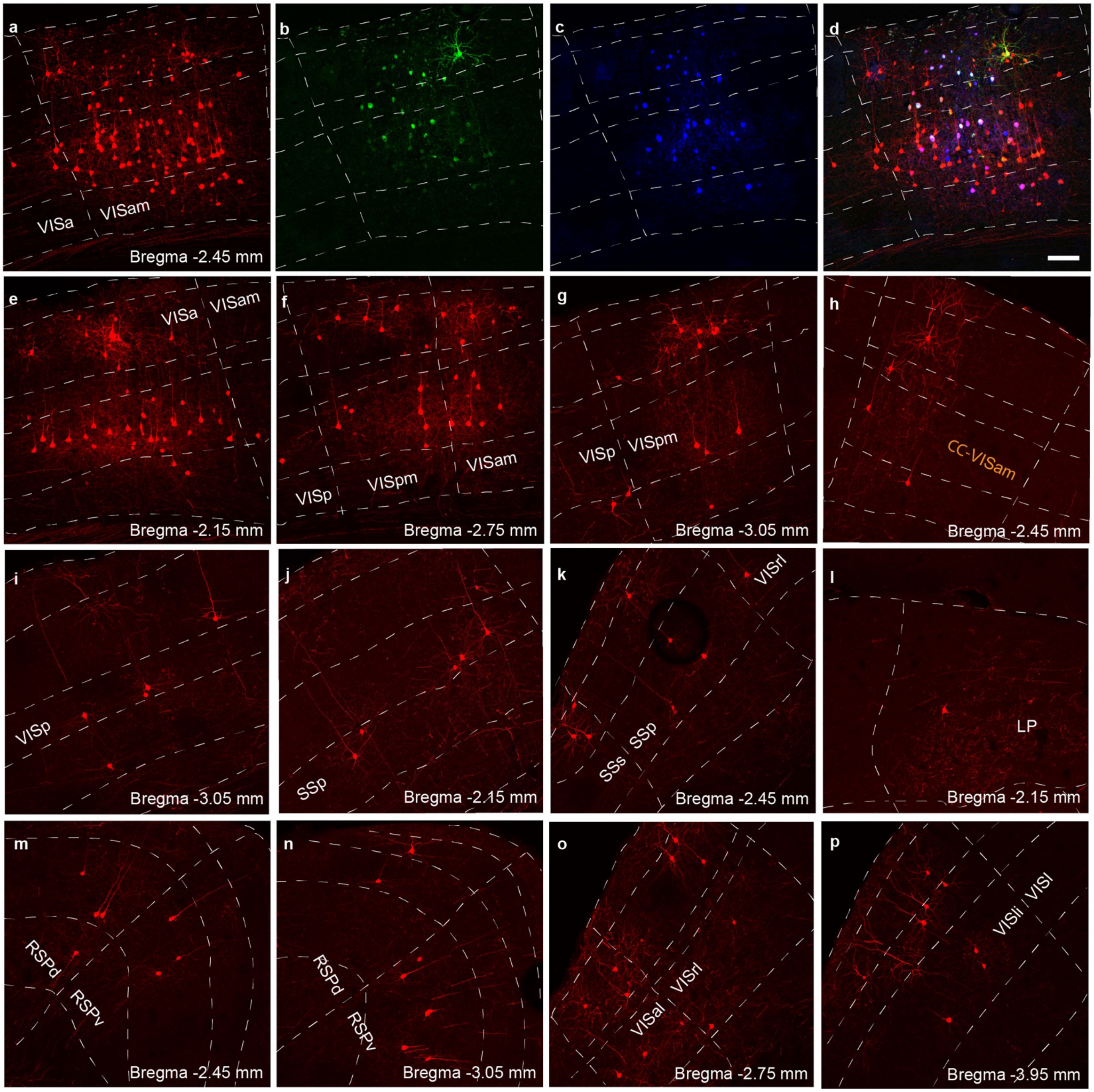


**Extended Data Fig. S16: Example confocal images of injection site and input regions in mouse used for longitudinal functional calcium imaging.** See Fig. 4a for diagram of experimental design. These images show that inclusion of the jGCaMP7s AAV did not prevent successful monosynaptic tracing. **a-d,** Injection site in V1. **a,** tdTomato, **b),** jCaMP7s, **c,** mTagBFP2, **d,** merge. Scale bar: 100 µm, applies to all panels. **e-p,** Inputs to parvalbumin-expressing V1 neurons are found in many different brain regions: other visual areas (**e-g**, **i**, **o-p**), contralateral cortex (**h**), somatosensory areas (**j-k**), thalamus (**l**), and retrosplenial areas (**m-n**). VISa, anterior visual area; VISam, anteromedial visual area; VISp: Primary visual area; VISpm: Posteromedial visual area; CC-VISam: Contralateral cortex - Anteromedial visual area; SSs: Supplemental somatosensory area; SSp: Primary somatosensory area; LP: Lateral posterior nucleus of the thalamus; RSPd: Retrosplenial area, dorsal part; RSPv: Retrosplenial area, ventral part; VISal: Anterolateral visual area; VISrl: Rostrolateral visual area; VISli: Lateointermediate visual area; VISl: Lateral visual area.

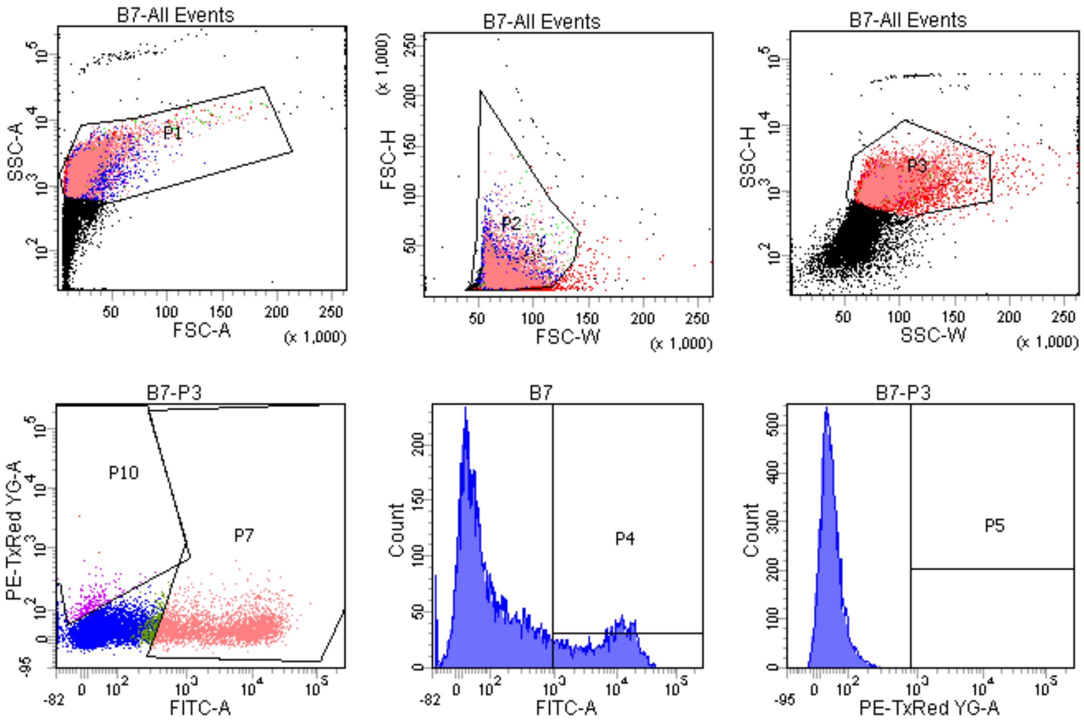


| | |
|---|---|
| Experiment Name: | 2020-11-24_dGL-4Cre and-4Fl... |
| Specimen Name: | RVdGL-4Flpo(EnvA)_III conc |
| Well Name: | B7 |
| Record Date: | Nov 24, 2020 1:07:31 PM |
| $OP: | HeatherSullivan |
| GUID: | 345bd03b-03fe-4027-b2f6-2b3... |

| Population | %Parent | FITC-A Mean | Pacific Blu... Mean | PE-TxRed ... Mean |
|---|---|---|---|---|
| P4 | 22.7 | 9,013 | 21 | 41 |
| P10 | 2.1 | 17 | 14 | 172 |
| P6 | 0.0 | #### | #### | #### |
| P8 | 0.0 | -20 | 124 | 3,393 |
| P5 | 0.0 | 6 | 68 | 2,116 |
| P9 | 33.0 | 6,338 | 18 | 40 |
| P7 | 30.4 | 6,871 | 20 | 39 |
| P11 | 0.0 | 6 | 68 | 2,116 |

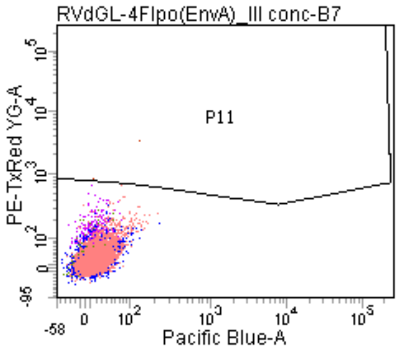


| Population | #Events | %Parent | %Total |
|---|---|---|---|
| All Events | 20,739 | #### | 100.0 |
| P1 | 11,313 | 54.5 | 54.5 |
| P2 | 10,064 | 89.0 | 48.5 |
| P3 | 10,000 | 99.4 | 48.2 |
| P4 | 2,266 | 22.7 | 10.9 |
| P5 | 2 | 0.0 | 0.0 |
| P6 | 0 | 0.0 | 0.0 |
| P8 | 1 | 0.0 | 0.0 |
| P9 | 3,304 | 33.0 | 15.9 |
| P1( | 214 | 2.1 | 1.0 |
| P7 | 3,036 | 30.4 | 14.6 |
| P1 | 2 | 0.0 | 0.0 |

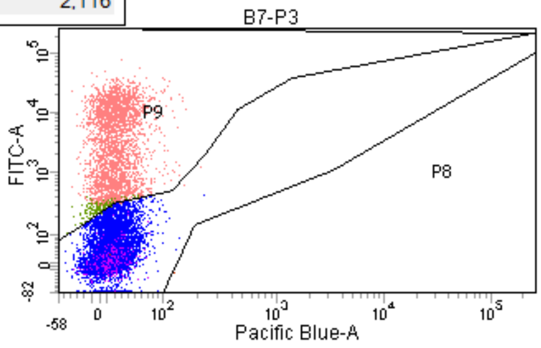




**Extended Data Fig. S17: Representative example of FACS data for titering viruses.** This example, included as stipulated by the Nature Portfolio Reporting Summary, shows FACS analysis data for one dilution of RVΔGL-Flpo(EnvA), used to infect HEK 293T cells which were immunostained for the rabies virus nucleoprotein using a blend of FITC-conjugated monoclonal antibodies (see Methods). The middle histogram in the second row shows the characteristic bimodal distribution (less distinct with second-generation RV vectors than with first-generation ones) with the uninfected cells in the mode on the left and infected ones in the one on the right. Gates are set by comparison with negative control wells of uninfected cells.
