## Supplementary figures and images for "Long-term labeling and imaging of synaptically-connected neuronal networks *in vivo* using double-deletion-mutant rabies viruses"

### Extended Data Fig. S4: Series of whole-brain images of labeled inputs to corticostriatal neurons using dG and dGL RV vectors expressing Cre

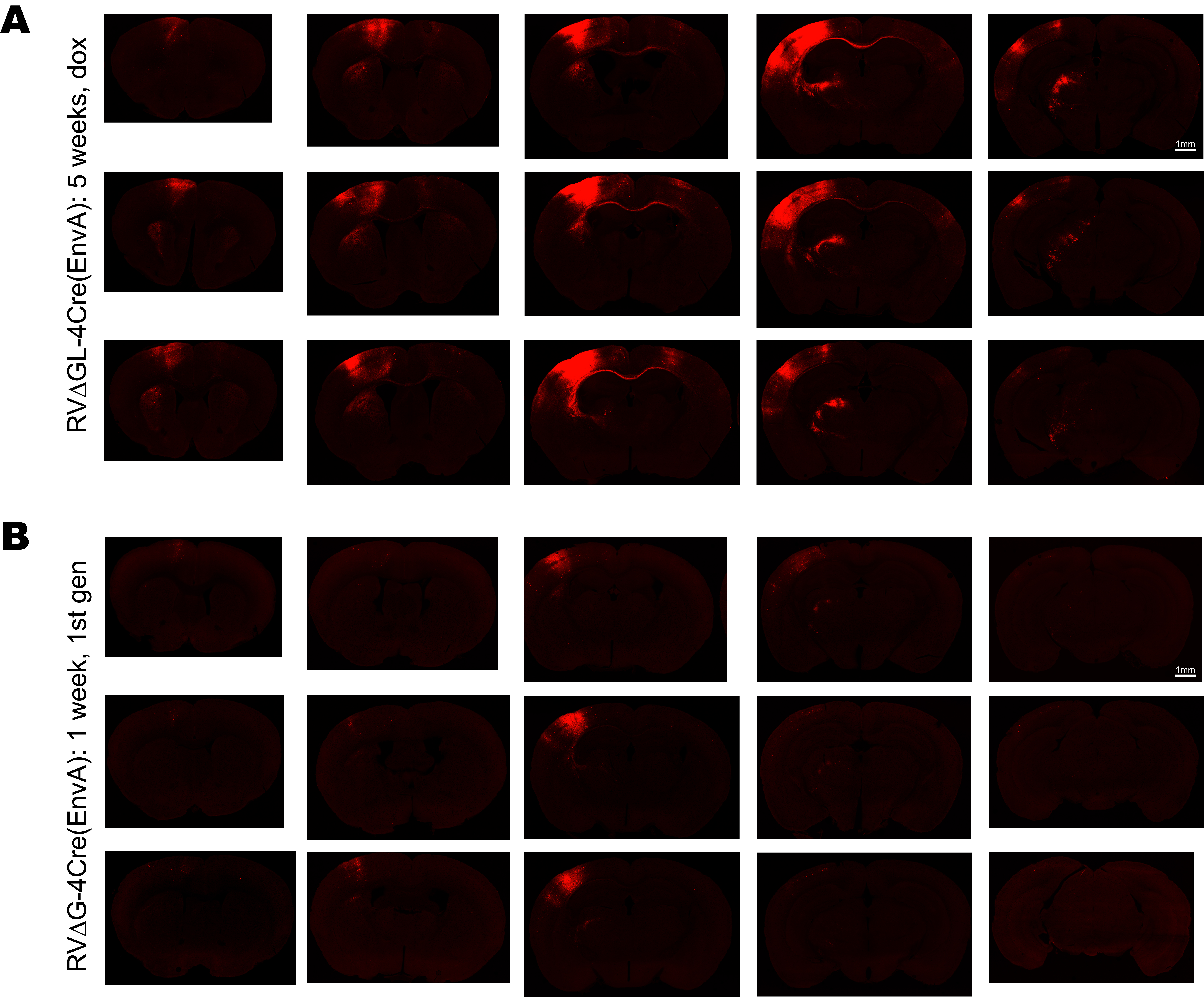

### Extended Data Fig. S7: Whole-hemisphere image series of labeled inputs to dopaminergic neurons using dG and dGL RV vectors expressing Cre

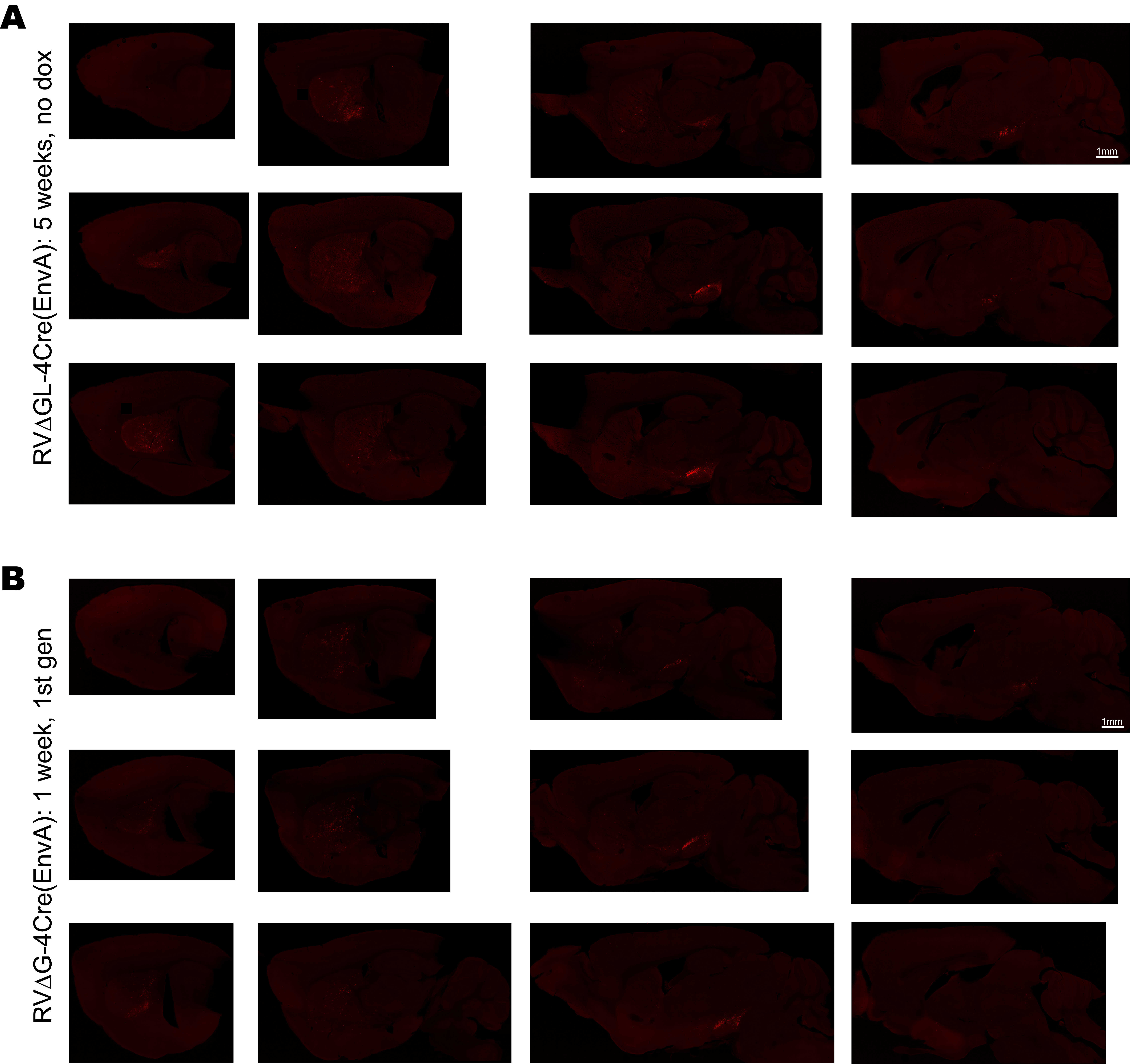
